# Inferring organ aging, hallmark of aging and senescence scores from blood and facial photographs

**DOI:** 10.64898/2026.09.23.753830

**Authors:** Kevin Schneider, Minja Belic, Matias Fuentealba, Fiona Senchyna, David Furman

## Abstract

Aging progresses asynchronously across organs, motivating the development of organ-resolved aging clocks. Direct assessment of organ aging in living humans, however, is largely impractical. Here, we construct tissue-specific transcriptomic aging clocks and infer organ biological age from paired whole-blood (blood-organ age) and tissue RNA sequencing data in the GTEx project. We observe high variability both in the strength of aging signatures across organs and in the ability to predict organ age and organ scores from blood. Pathway analysis shows that organ aging clocks are dominated by biological processes that differ across tissues. Age-associated multi-organ modules identified in an aging multi-organ network correlation analysis and organ age features causally mediated through organ-blood-organ transcriptomic interactions support a role for systemic signaling in coupling organ aging processes.

Given the established role of cellular senescence as a driver of tissue dysfunction and its tissue-specific accumulation during aging, we extend this framework to estimate burden of organ-level senescence and hallmarks of aging. Using senescence- and hallmark-associated gene sets, we derive tissue-specific senescence and hallmark scores (organ scores) and demonstrate that these burdens across multiple organs can be inferred from blood transcriptomes (blood-organ scores).

Finally, in pursuit of extending these models to other blood omics and alternative non-invasive assessments, we evaluate whether blood methylation (methylation-blood-organ age and scores) and facial photographs (facial-blood-organ age and scores) can serve as proxies for blood-organ age and blood-organ scores in independent cohorts (Edifice Health and Health and Retirement Study). The methylation- and facial-blood-organ age and scores were associated with mortality in additional testing cohorts (Framingham Heart Study and IMDB-WIKI dataset). These results establish a cross-modal framework for inferring organ-specific aging and senescence from minimally invasive data.

## Introduction

A major goal of aging research is to quantify biological aging more accurately than by chronological age. This has led to the development of molecular aging clocks, many of which robustly correlate with health outcomes such as mortality and multimorbidity^1^. Early epigenetic clocks demonstrated that DNA methylation patterns across tissues encode a strong aging signal^2^, a finding later extended to pan-mammalian clocks spanning multiple species^3^. Parallel efforts have produced transcriptomic^4^ and proteomic aging clocks^5^, reinforcing the concept that aging leaves measurable molecular imprints across biological systems.

Despite their predictive power, these system-wide clocks tend to lack clinical interpretability and intervenability. Accumulating evidence indicates that aging is heterogeneous both between individuals and across organs within the same individual^6^. Analysis of murine transcriptome has shown that while organs share a subset of conserved aging pathways, the magnitude and onset of age-associated changes are tissue-specific^7^. Consistent with this, pan-tissue aging estimators often exhibit differential accuracy or bias across organs^8^, motivating the development of tissue-specific aging clocks that better capture organ-level aging trajectories^4^.

Direct measurement of organ aging in living humans, however, is largely infeasible due to the ethical and technical limitations of tissue biopsies. To overcome this barrier, recent studies have leveraged circulating biomarkers in blood to infer organ-specific aging. Blood-based proteomic clocks, developed using large cohorts such as the UK Biobank, have demonstrated that protein signatures linked to specific organs can predict organ-level disease risk, health span, and mortality^9,10^. These approaches show that minimally invasive blood measurements can capture organ-resolved aging information at population scale.

Here, we propose a complementary strategy based on blood transcriptomics. Unlike prior blood proteomic studies, we leverage paired whole-blood and tissue transcriptomic data from the GTEx project^11^ to directly model blood–organ relationships. Transcriptomic profiles represent upstream readouts of regulatory programs and cellular state transitions and may therefore reflect different aspects of biological aging that bypass short-term compensatory regulation of protein abundance.

Previous work has shown that a substantial fraction of tissue-specific gene expression can be predicted from whole-blood transcriptomes^12^, supported in part by shared regulatory architecture and cross-tissue genetic effects. These observations suggest that blood transcriptomics can serve as a surrogate window into organ molecular states and enable inference of tissue-level aging programs without direct tissue sampling.

In this study we construct organ-specific transcriptomic aging clocks from tissue RNA sequencing data and use paired whole blood transcriptomes to derive minimally invasive predictors of organ biological age. In addition to chronological aging, we focus on cellular senescence, one of the key drivers of age-related organ dysfunction^13–15^, alongside other hallmarks of aging^7,16,17^. Senescent cells accumulate in a tissue-dependent manner during aging^18^ and secrete pro-inflammatory cytokines, chemokines, and matrix-remodeling enzymes, collectively known as the senescence-associated secretory phenotype (SASP), contributing to both local tissue degeneration and systemic aging phenotypes. We therefore extend our framework to estimate tissue-specific hallmark of aging scores and assess models to predict each from blood transcriptomes, providing a functional dimension to organ aging beyond aging acceleration alone.

Finally, these models were extended to blood methylation microarray and in pursuit of increasingly non-invasive aging assessment, we use facial photograph embeddings to predict blood-derived organ aging, hallmark of aging scores, and in particular senescence scores. Prior work has shown that facial images encode information about biological aging and health status^19,20^, suggesting that image-based proxies may capture aspects of systemic and organ-specific aging. We show that methylation-organ-age and facial-organ-age are associated with all-cause mortality in age-matched case/control testing cohorts. Our results establish an extensible cross-modal framework for inferring organ-specific aging and senescence from minimally invasive data and provide insight into the biological pathways that enable blood-based inference of tissue aging.

## Methods

This study used a multi-step approach to construct, test, and validate minimally invasive estimators of organ-specific age and hallmarks of aging scores. The data include whole blood RNA transcriptomics and microarray, DNA Methylation microarray and facial images (Figure 1). The workflow comprised the following steps for thoroughness and to identify the optimal model and gene set:

a. Splitting samples stratified by organ into training and test sets
b. Constructing organ-specific aging clocks with all genes (organ age all genes)
c. Constructing organ-specific aging clocks from genes filtered by those correlated with age (organ age genes correlated with age)
d. Constructing organ expression models from blood genes correlated with organ genes
e. Constructing organ-specific aging clocks from genes correlated with age and predictive through blood (organ age)
f. Constructing organ-specific aging clocks from blood transcriptome filtered by those correlated with predicted organ age (blood-organ age)
g. Calculate predicted organ age from organ expression predicted from blood using coefficients from (e)
h. Score organ samples using hallmark gene sets classified by correlation with age (organ score)
i. Construct hallmark scoring models from organ and blood transcriptomics (blood-organ score)
j. Construct organ-specific aging clocks using facial images in the Edifice test set (facial-blood-organ age and score)
k. Construct organ-specific aging clocks using DNA methylation in Health and Retirement Study test set (methylation-blood-organ age and score)
l. Test facial-blood-organ aging clocks on IMDB-WIKI image dataset
m. Test methylation-blood-organ aging clocks on Framingham Heart Study

**Figure 1.**
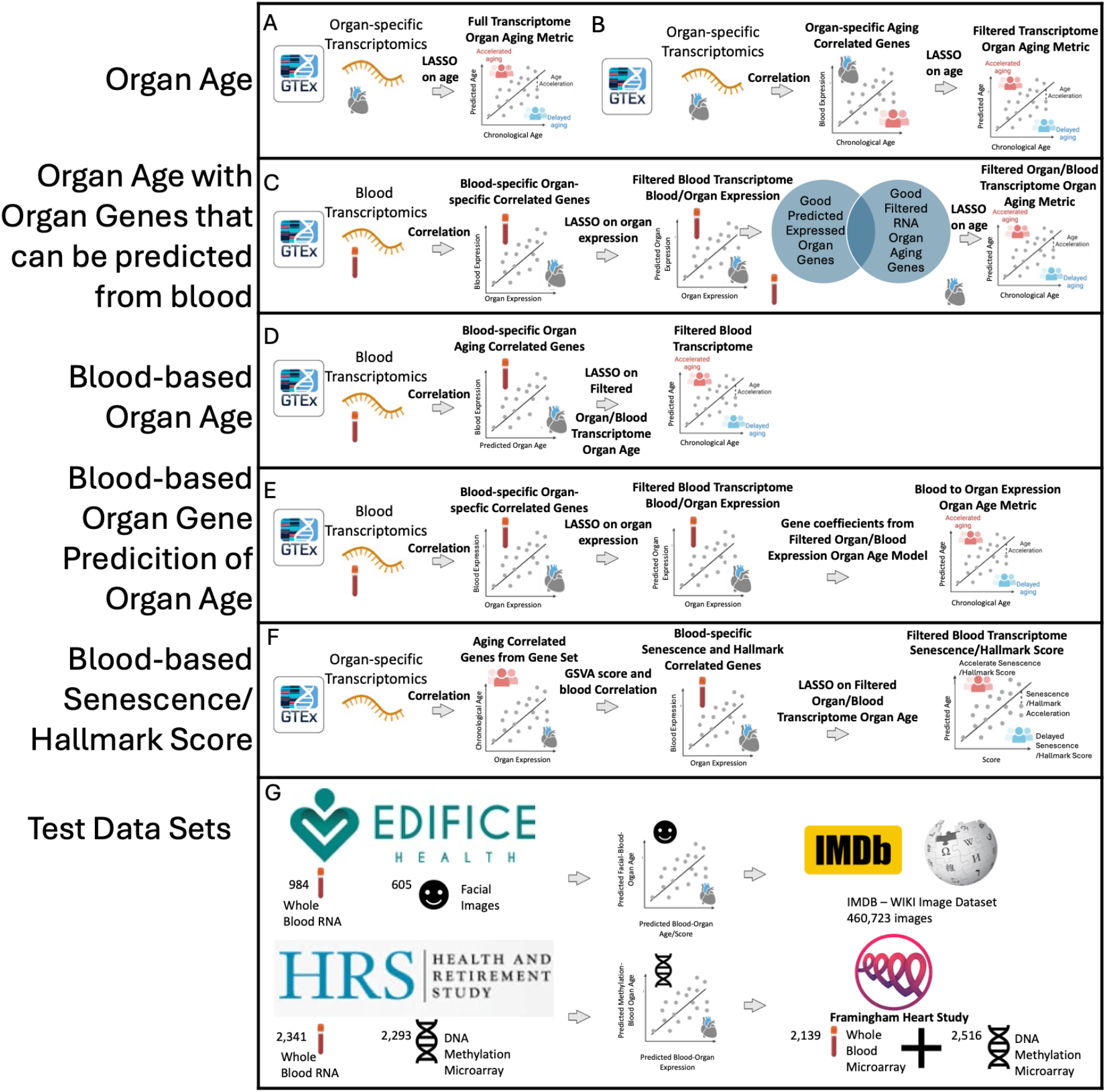
Characterization of Blood-Organ Correlates and Clocks. A) Organ clocks were derived from all expressed genes in each organ with over 20 individuals in GTEx. B) Organ clocks were derived using genes filtered by Pearson correlation with age. C) Organ clocks were derived using the genes from (B) and filtering for those that can be predicted from blood expression. D) Blood-organ clocks were derived from blood genes correlated with organ age from (C). E) Predicted organ expression from blood was used to calculate an alternative blood-organ age. F) Senescence and hallmark of aging gene sets were correlated with age and used to score organ expression. Blood expression was filtered with Pearson correlation with the organ score and used to construct a model for a blood-organ score. G) Blood-organ age was predicted in whole blood expression from the Edifice Health^20^ cohort and facial embeddings were filtered with Pearson correlations and used to construct a model of facial-blood-organ age that was predicted in the IMDB-WIKI^23^ image training set to associate with mortality. Blood-organ age was predicted in whole blood RNA from the Health and Retirement Study^21^ and methylation microarray data was used to construct a model of methylation-blood-organ age that was predicted in the Framingham Heart Study^22^ to associate with mortality.

Model fitting, hyperparameter tuning, and feature selection were performed using the same ten-fold cross-validation on the training set for all models (80/20 train/validation split). This cross-validated model was used for testing with the remaining test set that was not seen by the model and used for more unbiased testing within the same dataset. The same samples were used in the training set for all correlations and models in steps (a) through (i).

## Cohorts

### GTEx

Blood and organ gene expression was downloaded from the Genotype Gene Expression program GTEx V8f11. The program profiled bulk RNA-seq across up to 54 non-diseased organ biopsy sites from nearly 1,000 post-mortem donors. All organ aging, senescence scores and hallmark of aging scores and corresponding blood-based surrogates were trained on this dataset.

### HRS

Data were obtained from the Health and Retirement Study (HRS), an on-going study since 1995^21^. Normalized blood transcriptomic data was summed over each gene symbol of two to the power of the log counts and methylation data scaled from 0-100.

### FHS

The Framingham Heart Study is a multigenerational, longitudinal cohort initiated in 1948 that has followed participants for decades to study cardiovascular disease and aging-related traits^22^. The FHS study was used for external testing of blood-organ and methylation-blood-organ age and scores. Death was calculated as the years to death from the year the measurement was taken.

### IMDB-WIKI

The IMDB-Wikipedia image training dataset was compiled by Rothe *et. al.*^23^ for development of a convolutional neural network to predict age, providing a resource of images and metadata. Time to death for each image was calculated using the year of photo subtracted from the year of death extracted from Wikipedia.

### Edifice Health

Data were obtained from the Pre-Market Trial iAge® intervention trial conducted by Edifice Health (clinicaltrials.gov ID: NCT04983017). This decentralized, double-blind, randomized, placebo-controlled study enrolled healthy ambulatory adults to characterize iAge in a real-world wellness population. Participants underwent assessments including cytokine panels, physical function tests, blood pressure, HbA1C, microbiome profiling, bulk RNA sequencing, functional intrinsic capacity domain measures, standard clinical parameters, and multiple patient-reported outcomes. Facial photographs were collected for a subset of individuals taken by an iPad with non-standardized background and lighting. The Edifice cohort was used to construct facial photograph proxies of blood-organ age and hallmark of aging and senescence scores.

## Age Correlation and Blood-Organ Expression Models

For each organ, blood and organ raw count gene expression data were normalized using the varianceStabilizingTransformation (VST) function from the DESeq2^24^ package with a design matrix of ∼1 for batch correction with microarray data and by counts per million (cpm) using the edgeR^25^ package for comparison with other RNA sequencing data sets. To ensure interoperability of the linear models, blood samples from GTEx were filtered to retain only genes also present in Edifice Health and HRS as cpm and FHS as VST normalized counts. Analyses in GTEx were restricted to tissues with at least 20 blood-tissue sample pairs.

For building blood gene expression predictors of organ gene expression, the training set was first used to calculate Pearson correlation between each organ gene and all blood genes. Genes were filtered to include 500-5500 genes based on a starting value of abs(R)=0.4 and increasing or decreasing (R) by 0.01 until the number of correlated blood genes was within this range for each organ gene. Cross validated general linear models (CV GLM) were constructed using cv.glmnet from the glmnet package and an alpha of 0.1^26^. Organ genes with at least ten non-zero coefficients at the lambda that provides the minimum error were retained. Age correlation was calculated using Pearson correlation between cpm expression and chronological age. All predictions for expression, age and scores used coefficients based on the lambda that provides the minimum error (“lambda.min”) in the model.

## Biological Organ Age and Blood-Organ Age

Models were constructed to predict organ age using all organ genes (Figure 1A), organ genes that were correlated with age (Figure 1B), organ genes that were correlated with age and predictive using blood expression (Figure 1C). Blood-organ age models were constructed using either blood genes correlated with organ age (Figure 1D) or predicted organ expression from blood of genes with nonzero coefficients in the organ age model constructed with organ genes correlated with age and predictive using blood expression plus the intercept of the model (Figure 1E). Genes were filtered to between 500-5500 organ genes versus age and 100-5500 blood genes versus organ age using the same criteria as the Pearson correlations between blood and organ expression. All models were constructed using ten-fold CV GLM with an alpha of 0.1 of the training set.

## Senescence and Hallmark Organ Score and Blood-Organ Senescence and Hallmark Scores

Transcriptomes of each organ in the training set was also used to calculate the organ-specific hallmark scores using the gene set variation analysis from the GSVA^27^ package and gene lists from the twelve Hallmarks of Aging^17^. As we are also interested in senescence, senescence scores were calculated from the six previously curated gene sets: GO^28^ Cellular Senescence, Reactome^29^ Cellular Senescence, Reactome SASP, SenMayo^30^, Cell Age Up^31^, and the Buck SASP atlas^32^.

The Pearson correlation of these scores with age were used to determine if there was any age association between gene sets and the organ expression. To better score these gene sets in relation to age in each organ, the Pearson correlation was used to split each gene list into increasing or decreasing with age at R=0; R=-0.1;0.1 and R=-0.2,0.2 (Figure 1F). Composite scores for each sample were calculated as GSVA(R greater than cutoff) minus GSVA(R less than cutoff). The non-split and composite scores were correlated with gene expression and models were constructed in the same method as organ age, replacing age with the score in the model. The models for organ score and blood-organ score were constructed with a 10-fold CV GLM with an alpha=0.1 with the training set and using the same fold split as the organ age models.

## Metrics

Correlation was employed to identify significant relationships between different organ ages. We used Pearson correlation coefficient (R) and root mean squared error (RMSE) between predicted and observed values of the training and testing groups to quantify model performance. Significance was established following correction for multiple comparisons via Benjamini-Hochberg procedure, setting false discovery rate FDR to 0.05. Beta estimates were calculated using scaled feature values from the HRS dataset with logit (binomial) or linear (continuous) regression with glm or lm, respectively.

## Facial-Blood-Organ

To determine if organ and senescence clocks could be estimated from face features, we used blood transcriptome data with paired facial photographs from the Edifice Health trial^33^ to learn to estimate each score in that cohort from facial images (80/20 train/test split). For the present analysis, we have 605 participants who met the following criteria: (i) a frontal facial photograph captured using an iPad that was suitable for computer vision analysis, and (ii) available metadata and clinical assessments, (iii) if data for more than one visit was available only the earliest visit would be used. Images captured on iPad devices were converted from .HEIC to .png format on the command line using ImageMagick.

Images from the IMDB-WIKI database images collated for the convolutional neural network to determine facial age from facial images and chronological age^23^ were downloaded for embedding extraction. Metadata for IMDB-WIKI include date of birth, image taken date, gender, face location, face score, second face score, celebrity name, and celebrity id.

For EH and IMDB facial images, a computer vision autocropper was first used to crop the face out of each image. If a face could not be detected the image was not included in the analysis. Next, for EH photos backgrounds were removed from the object using a background remover implementation in torch vision (backgroundremover). Embeddings from ArcFace^34^, FaceNet^35^ and VGGFace^36^ were extracted using python. Embedding features were filtered to the top 100-500 based on Pearson correlation with blood-organ age or blood-organ score. Image embeddings were batch corrected with ComBat from the SVA^37^ package between IMDB and EH, using EH as the reference dataset. Edifice Health samples were split into an 80% training and 20% testing set. Downstream linear models (glmnet) were constructed using these pretrained-model derived image embeddings to predict previously calculated blood-organ age and blood-organ scores and derive a facial-blood-organ age and facial-blood-organ score.

IMDB facial images were filtered for those with a face.score >=2, with an age at photo that was less than the age at death, with a second.face score of NaN, an age of photo >= 20 and a time of death since photo of <= 90. Individuals with more than one photo were sorted by age at photo and only one photo of each individual was retained. Samples were split into train and test sets with an 80/20 split and a censor of twenty years was set. Survival modeling used years to event or censor from the retained photo, setting an event to a censor if it was recorded more than 20 years after the photo. Features were z-scaled and modeled against all-cause mortality using cox partial hazard multi-variate analysis with one hundred boot-straps (set with coxph.control from the survival package^38^) and including sex in the design matrix to construct a model to predict mortality risk. Median scores were used to split high and low risk individuals in the test set for downstream AUC scores using the timeROC package^39^ and Kaplan-Meier curves with the survival package.

## Methylation-Blood-Organ Age

All paired HRS samples with both whole blood RNA expression and whole blood methylation microarray were used. Samples were split into an 80% training and 20% testing set. RNA-methylation expression and methylation-RNA expression models were constructed using the same methods as those used to construct blood-organ expression models (see above). Blood-organ age and blood-organ scores were predicted in all samples. Methylation was filtered to the top 100-500 CpGs based on Pearson correlation (using “complete.obs” and removing samples with NA values in the retained CpGs) to blood-organ age and blood-organ score to derive models to predict a methylation-blood-organ age and methylation-blood-organ score. Predicted blood-organ age was scaled from 20-100 for methylation-blood-organ age models, as individuals in HRS are all greater than fifty years old.

All FHS samples with paired microarray expression and microarray methylation data were used in this analysis. For each sample a predicted blood-organ age, blood-organ score, methylation-blood-organ age and methylation-blood-organ score were calculated. Time to death since measurement was set in the same method as the IMDB-WIKI set.

## Aging Multi-Organ Correlation Analysis

For GTEx samples with blood, but no organ expression, the organ expression was predicted for all organ genes with good models that passed filtering. The combined observed and predicted organ and blood expression for all samples were combined into one data frame with the organ as a secondary feature label. Genes were subset based on a Pearson correlation with age above or equal to 0.2 and a variance of 1. A whole genome correlation network analysis (WGCNA)^40^ with power derived from a softpower analysis for a signed WGCNA was constructed. Limma^41^ was used to confirm significance of each module’s association with age. This Aging Multi-Organ expression network Correlation Analysis (AMOCA) provides insights into key multi-organ pathways involved in organ aging.

## Mediation analysis

Mediation analysis was completed between the log normalized count per million (CPM) gene expression in paired organs and blood. Only samples with observed organ and blood expression were used (predicted organ expression was not used for mediation). Target genes were derived from the organ age model constructed with genes correlated with age and predictive from blood expression and those that had a non-zero coefficient in the model with lambda derived from the minimum error. Mediator genes were derived from the blood-organ age model constructed with blood genes correlated with organ age. Source genes were filtered using increasing (or decreasing) Pearson correlation with the mediator to derive between 1000 and 4000 X->M->Y triples for each organ pair. Mediation was calculated using mediate from the mediation package^42^ and bootstrapped with 1000 Monte Carlo simulations in the forward X->M->Y and reverse Y->M->X direction. Those triples with a significant FDR average casual mediation effect (ACME) in the forward direction and p-value >0.05 in the reverse direction were retained as potential organ->blood->organ age gene mediation effects. Heatmaps aggregate all mediations, full and partial, and are visualized with pheatmap^43^.

## Compounds to alter organ clocks and senescent predictors

Gene-Compound Enrichment Analysis (GCEA) was performed using the GCEA package^44^ to identify compounds targeting the genes that are associated with organ-age and blood-organ age. Only the genes with non-zero coefficients in these models were targeted.

## Results

### Organ Clocks and Blood-Organ Clocks

Biological organ age clocks were derived for each organ that had paired organ and blood transcriptomes from at least twenty individuals in GTEx. Gene sets were filtered on 80% of individuals available for a given organ with testing completed on the remaining 20% to construct organ age models. Gene sets for each organ were either unfiltered (Figure 1A) or filtered to those that were correlated with age (see Methods; Figure 1B). To further filter genes used to derive a biological organ age and improve the likelihood of predicting organ age from blood transcriptomics, only organ genes that could be predicted from blood transcriptomics were retained (Figure 1C and Supplementary Figure S1). This gene set was used to derive a 10-fold cross-validated biological organ age. In the same training set the blood transcriptome was filtered to those genes correlated with the predicted organ age and used to derive blood-based biological organ age (blood-organ age; Figure 1D). For comparison to direct blood-organ age; organ predicted expression from blood was used to calculate biological organ age from the coefficients of the organ clock models (Figure 1E).

Biological organ age clocks were derived using these five methods for each of the 48 organs in GTEx (Figure 2A) with at least twenty individuals. The spinal cord cervical, amygdala and kidney cortex had no variation in the test set, which leaves 45 organs with at least some variation in the predictions on the test set. The frequency of gene overlap for biological organ age between organs using either organ or blood transcriptomics was low with more frequent overlap between organs using blood genes (Supplementary Figure 2). Gene ontology enrichment analysis of the Biological Process gene set found no overlap between organ gene sets and the top enriched terms, which is not surprising given most genes are unique to a single organ (Supplementary Figure 3). Enrichment of the blood genes used to predict blood-organ age have fewer organs enriched for gene sets but do show an overlap between enriched gene sets of adipose subcutaneous and testis and of prostate and lymphocyte cells. All organs for blood-organ age predictions had a Pearson correlation with age above 0.35 (Figure 2B) and root mean squared error (RMSE) between 8 and 13 years. For 42 organs, correlations with the test set were lower than the training set, which decreased to near zero for two organs and below 0.3 for eleven organs (Figure 2B and 2C and Supplementary Figure 3). RMSE values also increased in most organs, with small decreases in RMSE in ten organs (Figure 2C). The organ most correlated with the test sets include thyroid, breast mammary tissue, colon sigmoid, artery tibial, skin and esophagus with a Pearson correlation above 0.6 in the test set (Supplemental Figure 4). The 34 tissues with correlation above 0.3 in the test set show the most promise in deriving an organ biological clock from blood.

**Figure 2.**
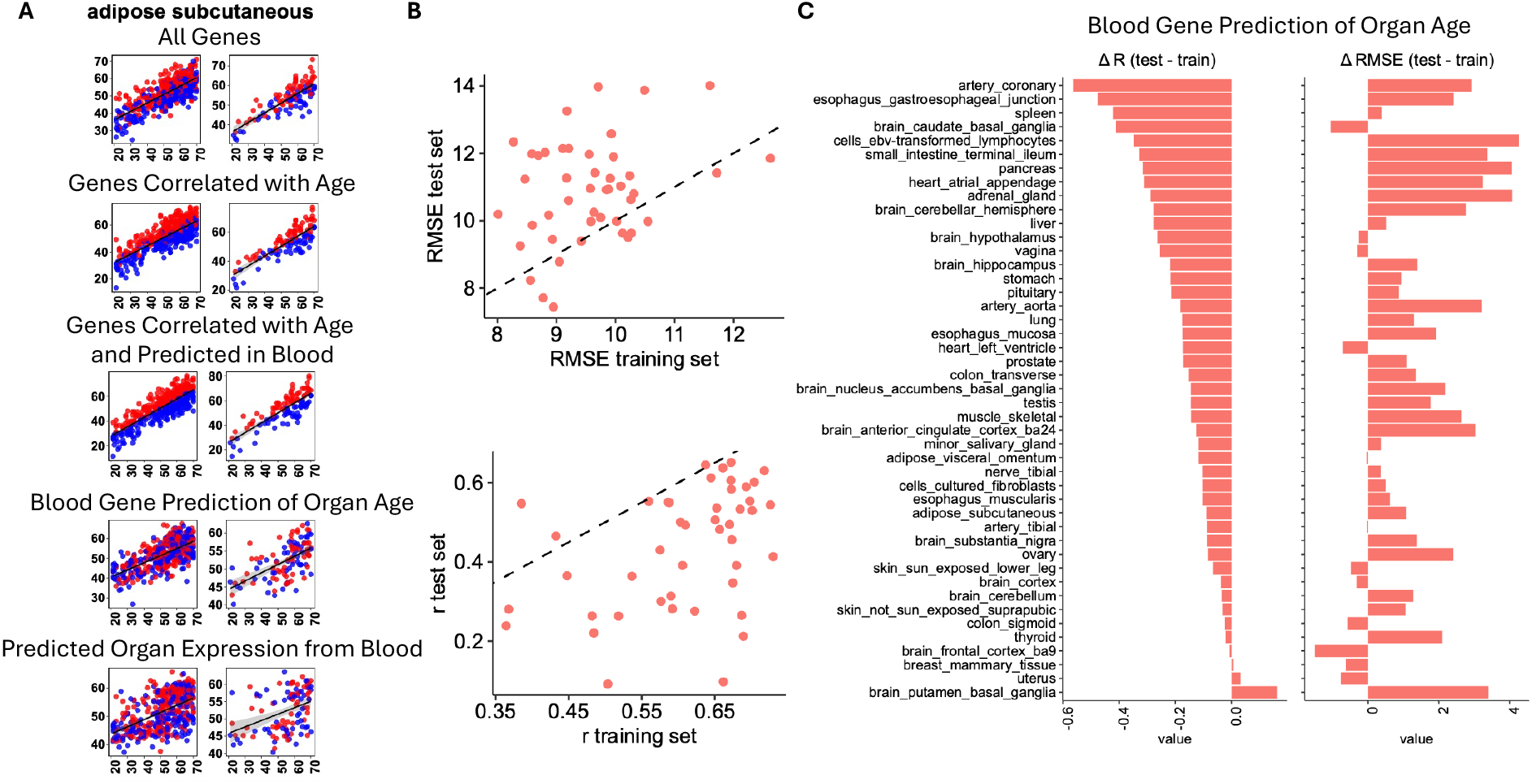
Blood-organ age models. A) The chronological age (x-axis) and predicted organ age (y-axis) is shown for one clock constructed using the methods detailed in Figure 1A – 1E for panels 1-5, respectively. Left panels are derived from the training set and right panels from the test set. Linear regression lines are shown as a black line. Colors are derived from the model detailed in Figure 1C and used for the construction of the Blood-organ age models. Blue dots have a negative predicted age residual and red dots have a positive predicted age residual with this model. B) For the 45 blood-organ age models with variance in the predicted test set, the predicted and observed values were used to calculate the RMSE and Pearson correlation between chronological age and the predicted value for the training and testing set. Dashed lines are shown for y=x. C) With the testing set Pearson correlations decrease with most blood-organ age models and RMSE values increase, but by no more than four. Some correlations go down near zero.

### Blood-Organ Gene Expression Correlates to Derive AMOCA Inter-organ Modules

In the training set, models were successfully constructed to predict organ expression from paired blood expression for about 35,000 to 45,000 genes per organ (Supplementary Figure 1) predicted from 26,369 blood genes. Blood genes with non-zero coefficients were shared by different organ gene models, with blood genes in 5 to 36,609 different organ gene models. Blood genes correlated with each organ gene (see Methods) were used to construct predictive models that were used to reconstruct the organ transcription from blood as well as derive the optimal genes to use for the organ models (see above). Organ expression predicted from blood was also used to derive a blood-based organ biological age using the coefficients from the organ-based model (Figure 1C). Although predicted organ age using this method correlates with age, the Pearson correlation with age is lower and RMSE is higher than directly using blood transcription to predict organ biological age. The reconstructed organ transcriptomes for individuals without sequencing data were combined with the actual organ transcriptomes from those with sequencing data. To perform an AMOCA that includes genes from all organs the gene set was filtered by variance above 1 and Pearson correlation with age above 0.2. Most modules identified contained a single organ for the genes contained within; however, three modules contain more than one organ of origin with module ME1 containing the most inter-organ correlation connections (Supplementary Figure S5).

### Causal Mediation of Organ Age through Blood Inter-organ Crosstalk

Mediation analysis was completed only with individuals that had available sequencing data. No reconstructed transcriptomics were included. Individuals with two paired organs and blood were used to estimate the casual effect of gene expression in a source organ (gene in organ X) on the target organ gene expression (gene in organ Y), for those genes used to predict biological organ age, as mediated through blood gene expression (gene M in blood), for those genes predictive of blood-organ age. Most mediations occurred locally from within the same organ (Figure 3A). A few organs, such as the nerve tibial, skin with sun exposure and testis showed casual association with multiple organs. Causal mediations are shown between organ X, gene M and organ Y for all organs (Figure 3B) and the detail the mediation of genes associated with adipose subcutaneous age (Figure 3C). This inter-organ cross-talk presents interesting targets to influence biological organ age via more distant organs.

**Figure 3.**
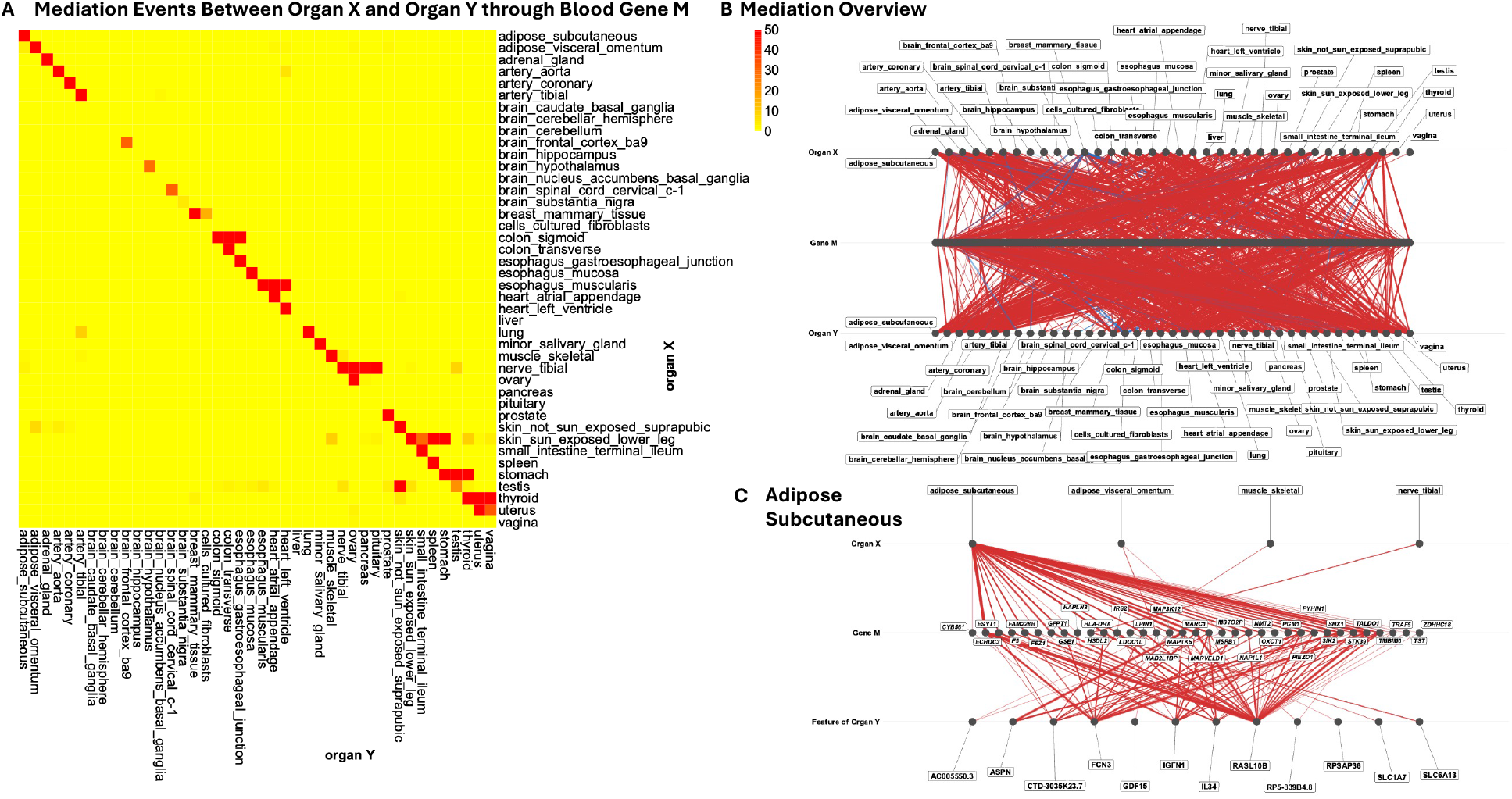
Inter-organ and intra-organ mediation of organ-age genes. A) Significant average causal mediation effects of genes from source organ X through blood organ M to target organ-age associated genes in organ Y. The total mediations through all gene triplets tested (X->M->Y) for each organ pair are shown (with a ceiling of 50 events). The most mediations occurred in a local manner within the organ between different organ-age associated blood genes. B) Topographical map of mediations between organ X through gene M to organ Y. Edge widths indicate the number of mediators in the edge. C) Organ-specific map showing mediations between organ X through gene M to genes in the adipose subcutaneous organ. The most mediations occur within the organ (organ X and organ Y are the same). RASL10B, a gene in the organ age model, is involved in the most mediations.

### Organ Aging Hallmarks and Senescence Enrichment

The aging hallmarks and senescence gene lists (see Methods) were used to score each sample in GTEx from all available organs and whole blood. These gene lists were derived from either large-language models^17^ or from published gene/protein lists^28–30,32,45^. The gene set variation analysis (GSVA^27^) scores had no correlation with age when the entire gene set was used (No split in Figure 4A). Gene sets were split in two sets based on positive and negative correlation with age at Pearson

**Figure 4.**
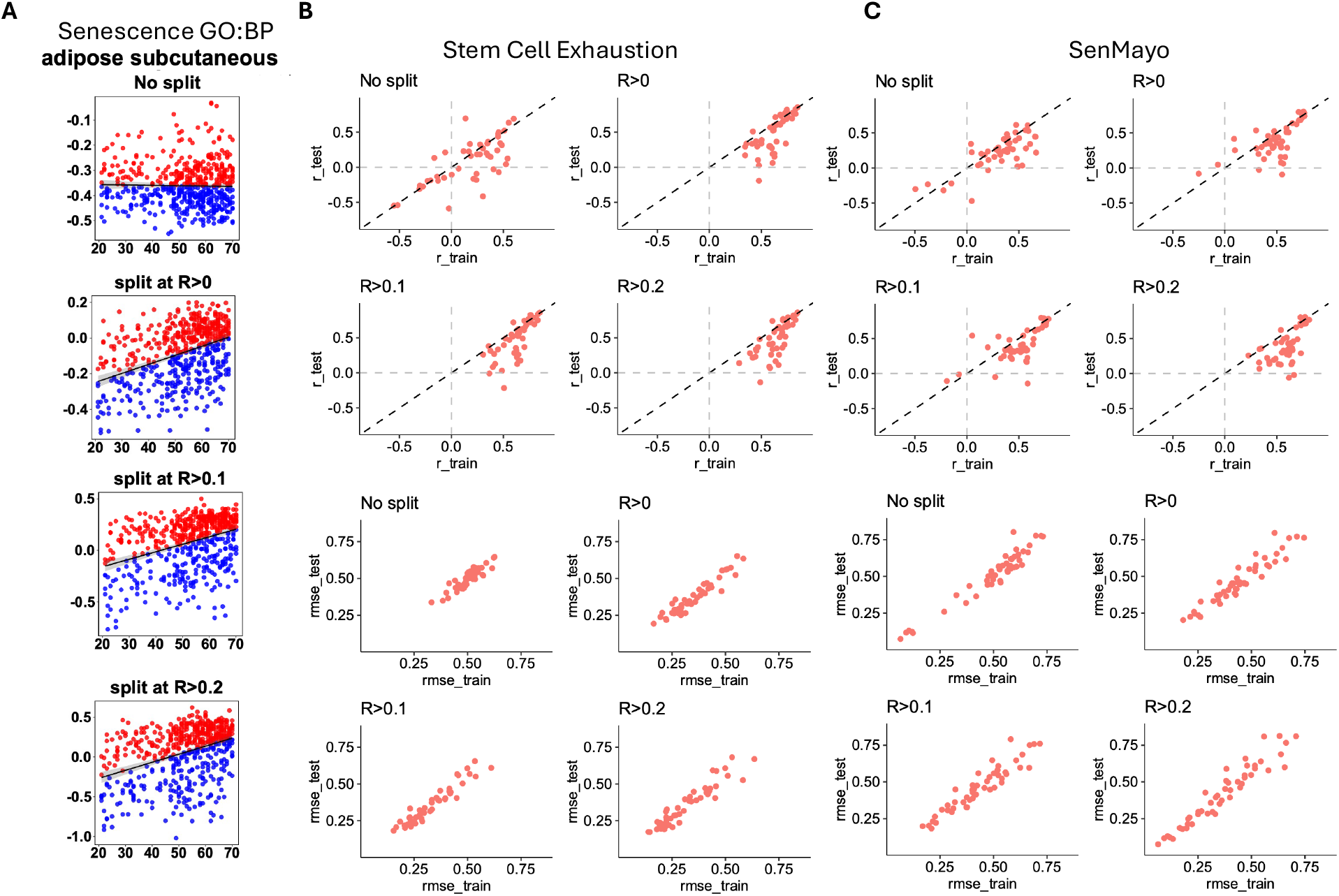
Senescence and hallmarks of aging scores and blood-organ score models. A) Organ expression was scored using GSVA for each gene set. Scores were compared with chronological age to determine if split gene sets based on Pearson correlation with age provide improved scores that increase and correlate with age. Splitting of gene sets, even at R=0, provide composite scores that increase with age. For the adipose subcutaneous organ, shown here, the scores increase in range as correlation requirements (R < -0.2 and R >0.2 range from -1 to 5 and R=0 range from -0.4 to 0.2). B) The Pearson correlation of the observed versus predicted values for the training and testing set of each blood-organ score model and each gene split for the Stem Cell Exhaustion hallmark of aging^17^ gene set are shown on top and RMSE on bottom. Pearson correlations are more frequently in the same direction after splitting of the gene set. RMSE values tend to be similar between the training and testing set. C) The results for the senescence gene set from SenMayo^30^ is shown. correlation of R=0 (R>0), R < -0.1 & R >0.1 (R>0.1), or R < -0.2 & R >0.2 (R>0.2). For each organ the split sets were scored independently for each sample. Composite scores of GSVA(positive gene set) + -1*GSVA(negative gene set) had a positive Pearson correlation with age and increasing minimum and maximum scores with increasing correlation requirements (Figure 4A). To construct a blood predictive model of each organ hallmark or senescence score we used similar methods as biological organ age replacing age with the GSVA score, filtering organ genes, filtering blood genes and predicting the GSVA score with blood genes. With no split in the gene list some organs had negative correlations with the observed score and splitting at any correlation requirement improved Pearson correlation with the observed score (Stem Cell Exhaustion Hallmark in Figure 4B and SenMayo Senescence in 4C). The training set performed better than the test set with some organs having a Pearson correlation under 0.2 with the test set. The RMSE values were less affected by splitting of the gene sets. These models were created for twenty-one gene sets derived from each organ.

### Heterogeneity of Organ Clocks

Blood derived biological organ age clocks were calculated in two independent test datasets, the Edifice Health (EH) cohort^33^ (984 samples from one or more visits) and the Health and Retirement Study (HRS) cohort^21,46,47^ (2,341) for 3,325 samples (from 2,720 individuals). Both cohorts had whole blood collected in RNA PAXgene tubes and blood-organ age and blood-organ hallmark/senescence scores were calculated for all samples. The Pearson correlations between different organ age residuals are mostly positive (Figure 5A), however certain pairs of organ clocks show an unexpected negative correlation or no meaningful correlation at all (Figure 5B). Visual examples of organ age depicted on anatomical plots colored by organ age were created and show how organ age acceleration and deceleration occur in one individual (Figure 5C).

**Figure 5.**
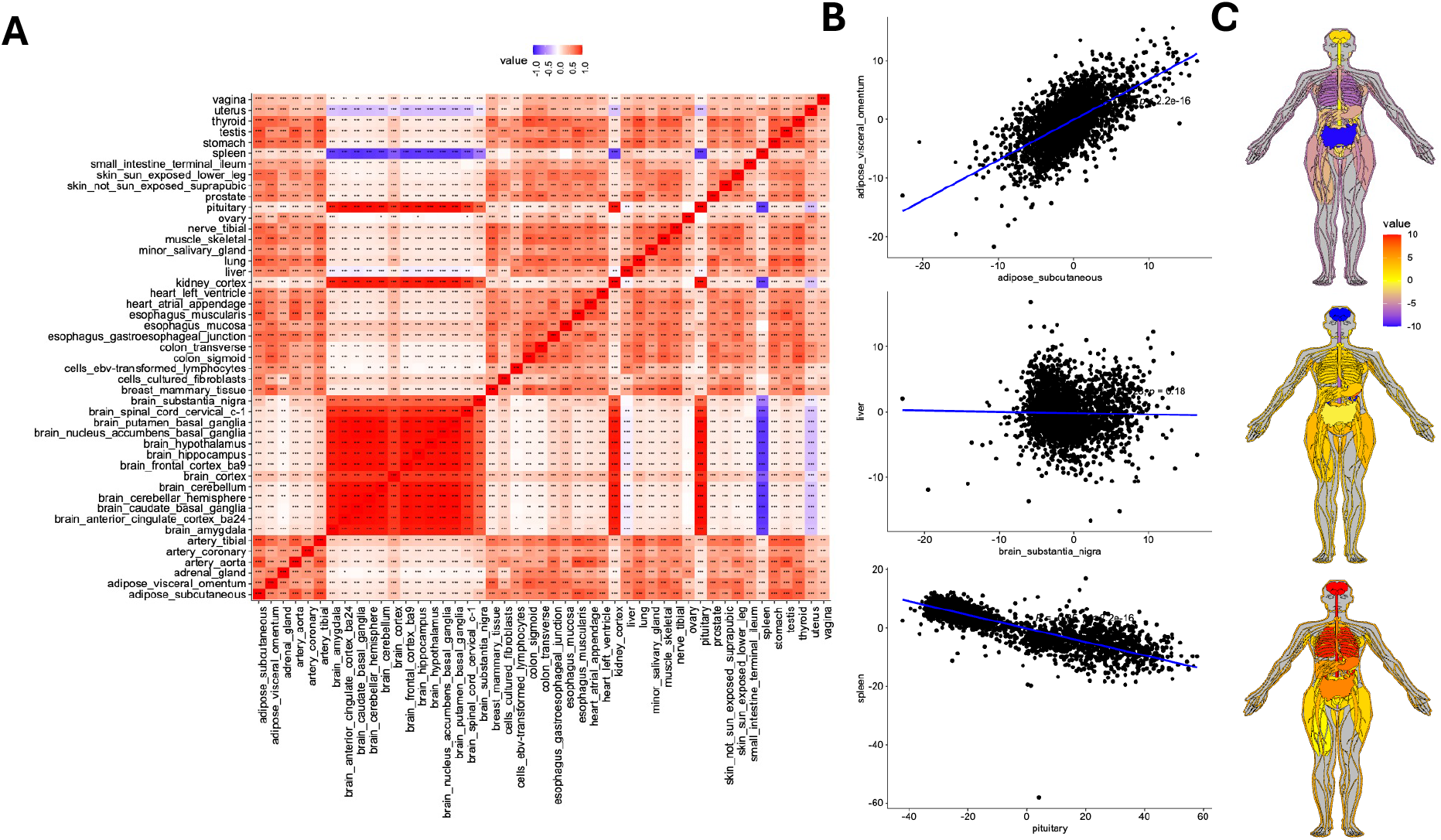
Systems wide organ age estimates in HRS and Edifice Health. A) Blood-organ age residual scores for 3,325 individuals derived from blood gene expression from the baseline visit. Pearson correlation of organ age scores between different organs are shown with most blood-organ ages positively correlated with each other. Some organs have ages that are negatively correlated or not correlated. B) Organ age scores for a positively-, negatively- and un-correlated blood-organ age pair. C) Visual example of organ age depicted on an anatogram plot and colored by organ age residuals for three individuals.

### Facial image Surrogates of Organ Age, Hallmarks and Senescence

Organ biological age models were constructed from facial image embeddings in the Edifice Health cohort that were correlated with the blood-organ age of 605 individuals, using only one visit if more than one visit was available (see Methods). Additional facial-RNA expression models were constructed for 26,488 different genes. Thirty-nine organs had a Pearson correlation above 0.2 for both blood-organ age and age (Figure 6A). There is an increase in the range of RMSE values up to 30 from blood-organ age and 40 from age. To test this model, faces from the IMDB-Wiki face images database^23^ with age, year at time of photo and face score were used with the extracted year of death to build a facial image mortality predictor from facial-blood-organ age output. Facial embeddings were extracted and images filtered using previously calculated quality metrics.

**Figure 6.**
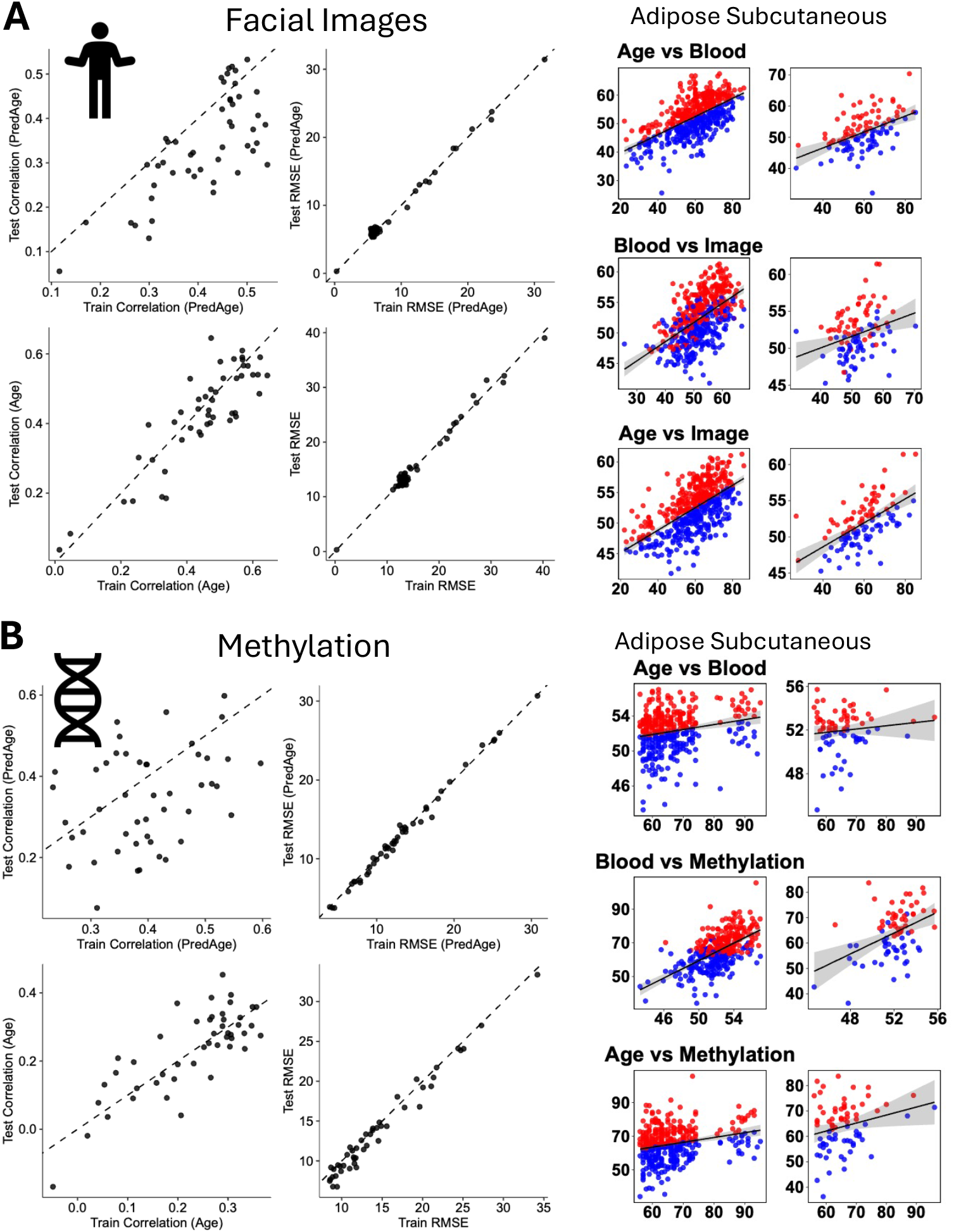
Facial-blood-organ and methylation-blood-organ age and score models. A) Facial image samples were split into training and testing sets to construct models as proxies for blood-organ age and blood-organ score and predict these in a subset not seen by the model in the Edifice Health cohort. Clockwise, black and white scatter plots show Pearson correlation with predicted blood-organ age, next are the RMSE versus predicted blood-organ age followed by age and then Pearson correlation with age. Overall Pearson correlation and RMSE scores for facial-blood-organ age is shown versus blood-organ age values predicted in this test cohort. RMSE values and correlations with age between training and testing were highly correlated. Scatter plots for samples from the adipose subcutaneous organ is shown at right. Samples are colored versus comparisons with facial output with the top and bottom chart colored by positive (red) and negative (blue) residuals and middle chart using the classifications from the top chart. B) Methylation microarray samples were split into training and testing sets to construct models as proxies for blood-organ age and blood-organ score and predict in a subset not seen by the model in HRS. Black and white scatter plots are in the same order as (A). For methylation-blood-organ age the overall Pearson correlation with blood-organ age was more varied, with more models having a R <0.2, which is also seen with age. Samples are colored versus comparisons with methylation output with the top and bottom chart colored by positive (red) and negative (blue) residuals and middle chart using the classifications from the top chart.

Images were filtered to remove images with missing information, odd calculations (negative or very high age values at time of photo), and a low facial quality score. The earliest image was used for individuals with more than one image and were matched one-to-one for age between alive and dead and split into an 80/20 train/test set. A multivariate cox partial hazard analysis identified the age of the left ventricle of the heart, colon transverse, ovary, putamen basal ganglia, breast mammary tissue, minor salivary gland and vagina as increasing risk of all-cause mortality in all individuals (Supplementary Figure S6A), the left ventricle of the heart, colon transverse, esophagus gastroesophageal junction and putamen basal ganglia in males (Supplementary Figure S6B). No significant organs in females were identified (Supplementary Figure S6C) with this filtered cohort. Features with a positive hazard ratio were used to stratify individuals in high- and low-risk groups. The model with all individuals has an AUC score of 0.74 to rank an individual for twenty-year survival, with a lower AUC using sex-specific models.

### Methylation Surrogates of Organ Age, Hallmarks and Senescence

Organ biological age models were constructed from methylation microarray data available in the Health and Retirement cohort for 2,293 individuals with paired RNA sequencing data. CpGs shared between HRS and Framingham Heart Study (FHS) were correlated with blood-organ age and blood-organ hallmark and senescence scores and used to construct models to predict the blood-organ age and scores (see Methods). There are 31,955 blood transcriptomic genes predictive of 256,454 CpGs (30,615 were present in two or more CpG models) and 230,485 CpGs predictive of 28,330 blood genes (230,346 were present in two or more RNA models). The methylation-blood-organ age for thirty-eight organs had a Pearson correlation above 0.2 for blood-organ age and twenty-eight organs for age (Figure 6B). There is an increase in the range of RMSE values up to 30 from blood-organ age and age. Clinical blood measurements from HRS were associated with organ age scores with eosinophil counts and monocyte counts showing discordant associations between different organ age scores (Supplementary Figure S7). However, STNFR-1, IL-1RA, IL-10, IL-6, CRP and Cystatin C are consistently associated with higher organ age across many organ systems and decreasing lymphocyte and albumin levels are associated with lower organ age scores.

To test this model in an outside dataset we used the methylation and RNA microarray from the Framingham Heart Study^22^ (FHS) to predict blood-organ and methylation-blood-organ age. The FHS has a similar platform for methylation data as HRS, but their RNA platform used microarray technology (probe based on a slide) versus sequencing technology (amplified and read by a sequencer). Predicted blood-organ age and methylation-blood-organ age have positive Pearson correlations with age and each other (Supplementary Figure S8A); however, many organs show higher correlation and lower RMSE with age using methylation than RNA with this cohort (Supplementary Figure S8B). This is likely due to both the different technology platform and the fewer number of genes in that platform that can be used in the model. This is further exaggerated when comparing the methylation scores with the RNA scores (Supplementary Figure S8C).

To determine if the predicted methylation-blood-organ age scores are associated with mortality, a multivariate cox partial hazard analysis identified the age of the lung, kidney cortex, testis and small intestine terminal ileum as increasing risk of all-cause mortality in all individuals (Figure 7), the lung and brain cerebellum in males (Supplementary Figure S9A) and minor salivary gland in females (Supplementary Figure S9B). The model of mortality due to cardiovascular disease in all individuals (Supplementary Figure S9C) has the highest AUC score of 0.76 of the cohorts tested to rank an individual for ten-year survival, with a lower AUC using sex-specific models (Supplementary figure S9D).

**Figure 7.**
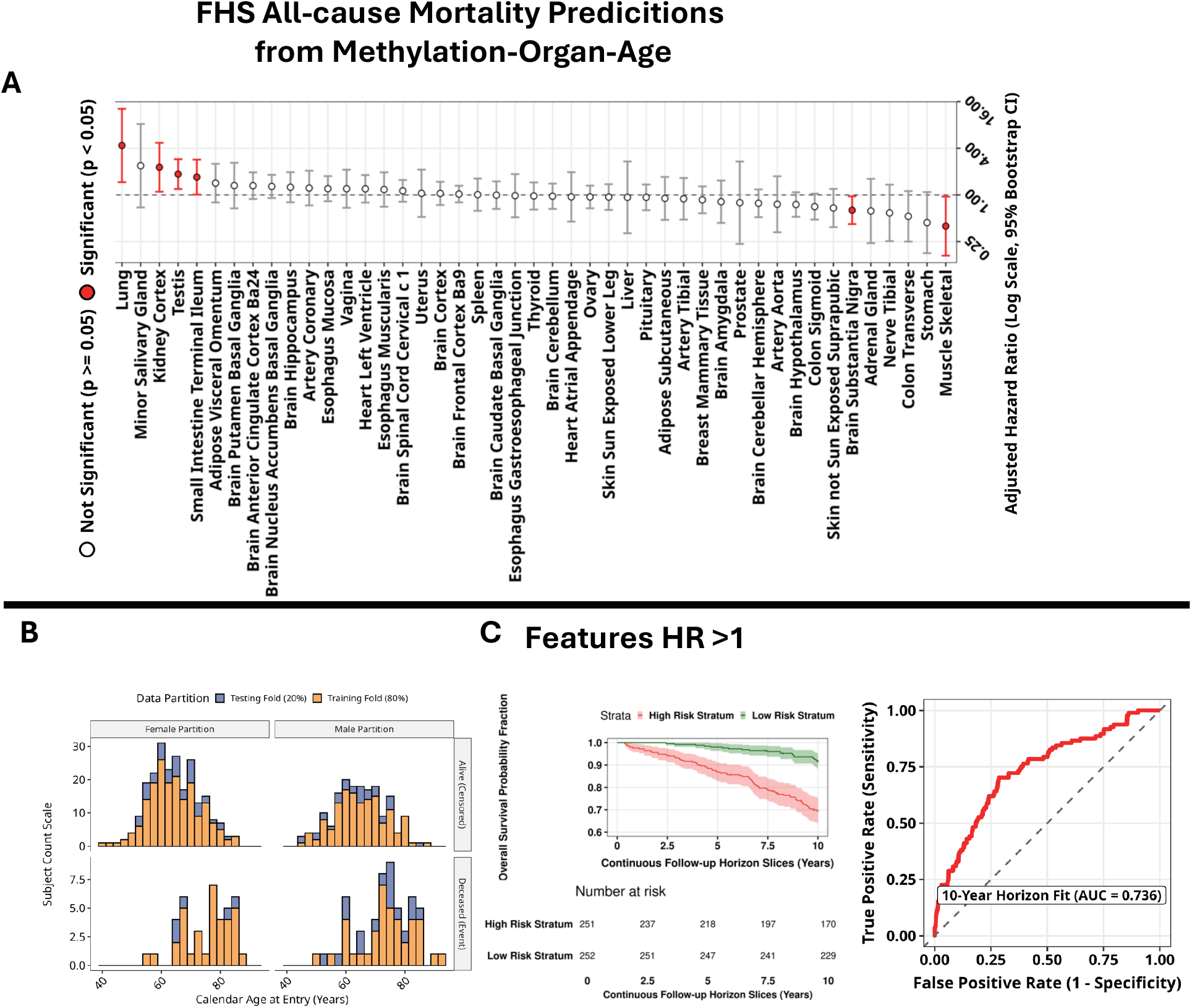
Methylation-blood-organ age predicts mortality in FHS. The methylation-blood-organ age was predicted in FHS and used to construct mortality multivariate cox partial hazard models. Groups were split into 80% training and 20% testing with models built on training data and survival plots and AUC calculated from the test set. A) Cox multivariate analysis identified organ age scores associated with survival. B) Population age and survival details on cohort used for cox analysis. C) Using only features with a positive hazard ratio from the cox model, the population was stratified by high and low risk. When adjusting for the confounding effects of each organ age, some organ age scores show protection as they increase. These nuanced features are more difficult to interpret and not included in the final model. The ten-year mortality Kaplan Meier curve and a time-dependent receiver under the curve (ROC) to rank an individual for ten-year survival was used to calculate the area under the curve (AUC).

### Compounds that Target Organ Age and Blood-organ Age

Gene compound enrichment analysis identified compounds that may alter organ genes and blood genes in each biological organ age and blood-organ age model. Compounds from DrugBank, FooDB and LINCSL1000 with nominal significance (P <.05) in the fisher overlap test (Supplementary Figure S10) are found to target one or more organs. The 20 compounds that alter the most organ systems from the organ and blood gene sets provide potential translational targets for improving organ age (Supplementary Figure S10). These compounds represent potential therapeutic targets associated with each organ clock.

## Discussion

This work centers on exploring minimally invasive ways of estimating aging and hallmarks of aging and senescence scores for individual organs. Specifically, we constructed organ-specific transcriptomic aging and hallmark of aging and senescence scores. Aging-correlated gene sets allowed calculation of senescent and hallmark scores that were shown to increase in each organ with age, versus scoring unsplit gene sets. The organ transcriptomes and the organ age and scores were inferred using paired blood transcriptomes. Models were tested on outside datasets and surrogates of blood-organ age and scores were constructed using facial images and blood methylation microarray data. The facial-blood-organ and score models were tested in the IMDB-WIKI image repository^23^. The blood-organ age and score models were tested with RNA microarray data and paired methylation-blood-organ age in FHS. The surrogates of these blood-organ age and scores provide valuable low-cost indicators of potentially high organ-specific biological age and high organ-specific senescence and hallmark of aging scores that is an indicator of mortality within ten years. The organ-specific biological ages show varied concordance with chronological age across different organs, partly due to varying sample sizes for different tissues, though it is also plausible that certain tissues show a weaker transcriptomic aging signature, such as the coronary artery and esophagus junction. Out of the modelled 45 tissues, 34 show high correlation between biological and chronological age (Figure 8). Prediction of organ age from omics and clinical markers have been shown to be associated with mortality and morbidity placing those omics and markers into specific systems for age calculation^48^ or using ICD diagnosis to associate those markers with the organ system^49^. Organ age predictions using GTEx expression profiles to identify blood proteins have also been used to build organ-system specific clocks associated with age and mortality^9,10,50^, but proteomics and transcriptomics are not derived from paired individuals and may or may not represent the blood-proteomic composition from that organ. Organ age residuals have also been modeled from histological slides paired with blood transcriptomics in GTEx^51^. This study directly correlates blood expression with organ expression in the same individuals giving organ age calculations that are directly connected to the organ of interest to offer direct insight into these associations. Clinical and cognitive features from HRS were correlated with each organ age score. Methylation-blood-organ age, from FHS, and facial-blood-organ age, from WIKI-IMDB dataset, were predictive of ten- and twenty-year mortality, respectively. In this multivariate analysis, some organ age scores are protective as they increase, which may be due to confounding effects between different organ age scores. The younger organs not associated with survival are more difficult to interpret and require further investigation into which organs may be causing this confounding effect. This study also allowed the analysis of the direct causal mediation of genes associated with organ age by other organ systems through blood. Mediation analysis identified RASL10B in adipose tissue as being involved in many mediations. This gene is known to be involved in the circadian rhythm in humans and baboons^52^ of white adipose tissue.

**Figure 8.**
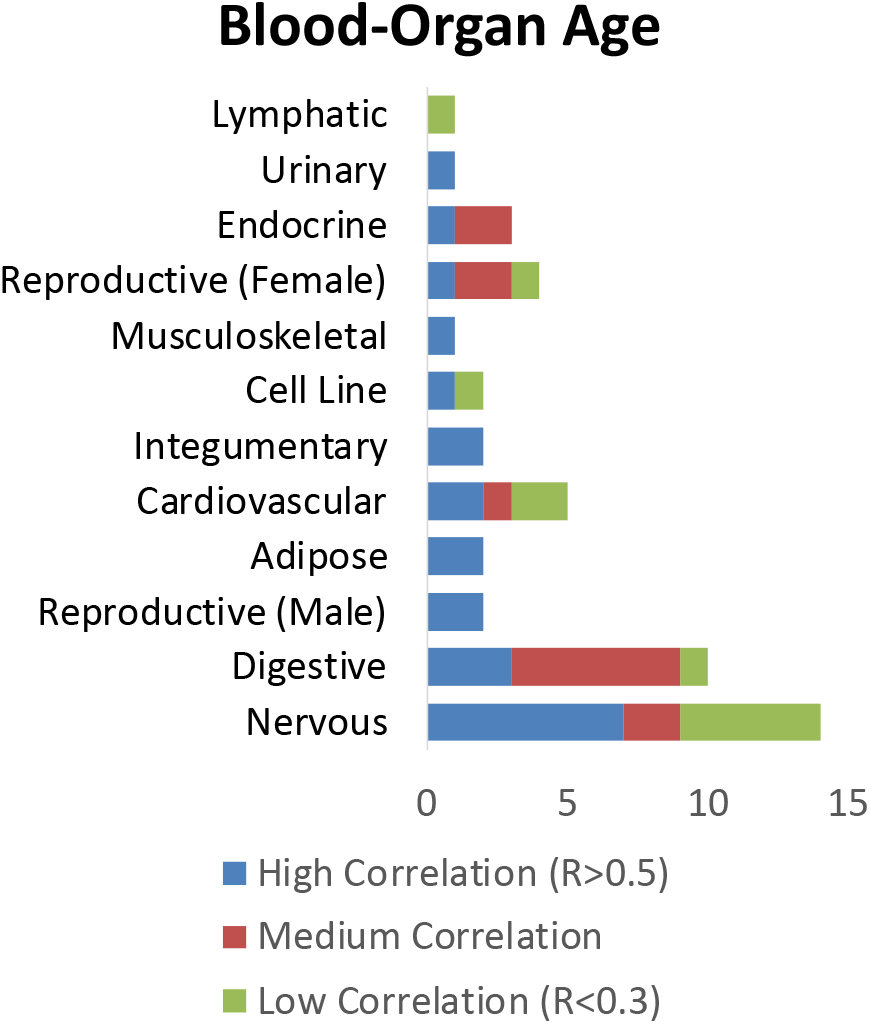
Organ systems with high Pearson correlation between chronological and blood-organ age in the testing set. Organs in the nervous and digestive system are most represented.

For the organ systems the nervous and digestive system were most overrepresented in the number of organs with models that correlated well with age (R>0.5), but many organs also showed weaker correspondence (mean R < 0.30). Across all organs, in the testing set pairwise correlations among organ age residuals (ΔAge) averaged r = 0.44 ± 0.15, indicating moderate systemic synchrony but substantial organ-specific deviation. These results demonstrate that transcriptomic variation in blood contains sufficient information to approximate organ-level biological aging states derived from organ transcriptomes, but with varied ability to predict organ age among tissues. This suggests variable coupling between tissue-intrinsic aging processes and systemic transcriptomic signals accessible in blood. Peripheral, visceral and glandular **t**issues such as thyroid, breast, colon, artery skin and tibial artery, colon sigmoid, esophagus muscularis and were highly mutually correlated (R >0.6 on the test set), consistent with coordinated vascular-metabolic aging mechanisms^53,54^. These organ models show a similar performance to those reported in RNAAgeCalc^4^. These results show that transcriptomic variation in the blood contains sufficient information to approximate organ-level biological aging states derived from organ transcriptomes. The highest correlation with chronological age were in peripheral, visceral and glandular and some of the lowest in the nervous system, suggesting that these on average show most age-related changes. Most tissues showing the lowest correlation with chronological age have slow cell turnover except for spleen and small intestines^55,56^.

This study, which relies on cross-sectional GTEx transcriptomes with variable sample sizes (20–400 per organ) and post-mortem intervals, has several limitations that should be addressed in future research. The sample size may not fully capture inter-individual variability, and larger cohorts could provide more robust and generalizable estimates. Additionally, cross-validation metrics may be overly optimistic due to potential biases associated with 10-fold cross-validation, highlighting the need for external validation using additional independent datasets. Finally, unmeasured confounding factors, such as lifestyle or environmental exposures, may influence gene expression patterns, and future studies should aim to account for and control these variables.

In line with work presented by Basu *et*. *al*.^12^ with TeeBot, we find that large proportions of gene expression in tissues can be estimated from blood transcriptome. The number and strength of blood–organ gene correlations varied widely by tissue. Across all organs, 26,369 blood genes were significantly correlated with at least one organ transcriptome. Blood-organ age calculated using blood-organ expression models does not perform as well as using blood expression correlated with blood-organ age. Organs within the same physiological system often shared overlapping correlated gene sets -for instance, digestive or respiratory tissues showed common epithelial and vascular modules-suggesting partially shared transcriptional channels within organ systems.

While blood-organ age and scores offer promising tools for assessing biological aging, their full potential can only be realized through rigorous validation and integration with additional types of data. Our findings have significant implications for personalized medicine by enabling non-invasive estimation of tissue-specific aging states using blood samples alone [7]. This approach could facilitate early detection of accelerated aging in critical organs such as the heart or kidneys before symptoms manifest clinically. Facial-blood-organ age and scores provide additional low-cost entries to enable individuals to identify possible high associated organ age or scores, which can be tested with more in-depth transcriptomics to identify interventions using GCEA. Quercetin was previously identified as a compound that could reverse aging signatures associated with simulated microgravity in peripheral blood mononuclear cells^44^. The compounds identified in this study targeting genes from both organ and blood-organ age and score models provide multiple potential targets to test *in-vivo* organoid models and future pre-clinical and clinical testing. Identification of drugs that have already completed a clinical trial may provide a more expedited path to being used. N-Acetyl-D-Glucosamine (also known as NAG) was identified as targeting the second highest number of organ age associated genes and has been used in previous clinical trials related to facial hyperpigmentation^57^ and shown in a small clinical trial of 34 multiple sclerosis patients to inhibit several inflammatory cytokines when taken orally^58^. Integrating multi-modal data will likely improve model accuracy and provide deeper insights into tissue-specific aging mechanisms. Longitudinal and multi-omics studies will be essential to determine whether changes in blood-derived organ age precede or mirror true physiological decline. Integrating proteomic, metabolomic, and methylation data could refine prediction accuracy and uncover additional causal mediators. Single-cell and spatial transcriptomics may identify cell populations responsible for cross-organ signaling, and large population cohorts could reveal demographic and environmental modifiers of blood–organ coupling.

This integrated multi-organ transcriptome study constructed models to predict organ age, organ senescence scores, organ hallmark of aging scores and organ expression from either blood transcriptomics, blood methylation microarray, facial images and to a lesser extent blood RNA microarray data. The organ age surrogates have shown how blood correlates can be used to predict many organ-specific features, which can be predictive of mortality and provide cheap multi-organ health assessment capabilities. We identify potential therapeutic compounds that can be personalized to target those organs with the highest age or mortality risk. We further developed models to derive blood RNA expression from blood DNA methylation, blood DNA methylation from blood RNA expression and blood RNA expression from facial images to provide additional biological information beyond just organ age and scores using the blood-organ expression models derived in this study. Inter-organ cross-talk mediated through blood and multi-organ modules identified with AMOCA provide function insights and potential targets for future studies of organ health-span.

## Acknowledgements

We would like to thank Edifice Health for providing data from their clinical trial. This research was supported by NIH grants U54AG075932, R03OD036497, U01-AG086214 and P01-AG066591. The Genotype-Tissue Expression (GTEx) Project was supported by the Common Fund of the Office of the Director of the National Institutes of Health, and by NCI, NHGRI, NHLBI, NIDA, NIMH, and NINDS. The data used for the analyses described in this manuscript were obtained from: Edifice Health, HRS, FHS and version 8 of the metadata and transcriptomic data from the GTEx Portal.

## Conflict of Interests Statement

D.F. and K.S. are employed by Edifice Health Inc. Edifice Health Inc. does not own any intellectual property which may arise from this study.

## Notes

### Competing Interest Statement

D.F. and K.L.S., are employed by Edifice Health Inc. Edifice Health Inc. does not own any intellectual property which may arise from this study.

