## Supplementary S1-S10 for "Inferring organ aging, hallmark of aging and senescence scores from blood and facial photographs"

**Supplemental Figures.**

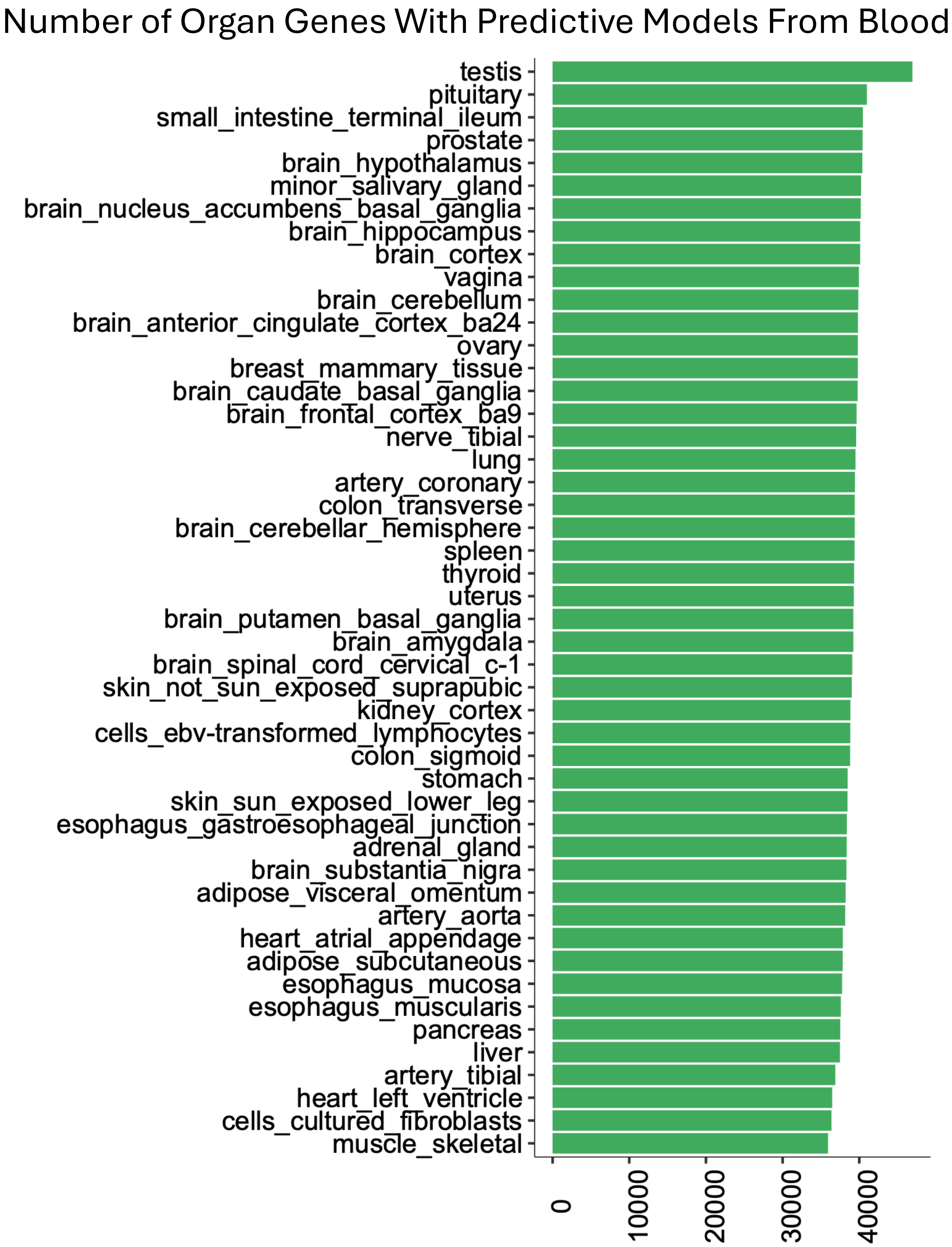


**Supplementary Figure S1. Blood expression models of organ expression.** Models for each organ gene were constructed and retained with at least ten non-zero coefficients at a lambda with minimum error.

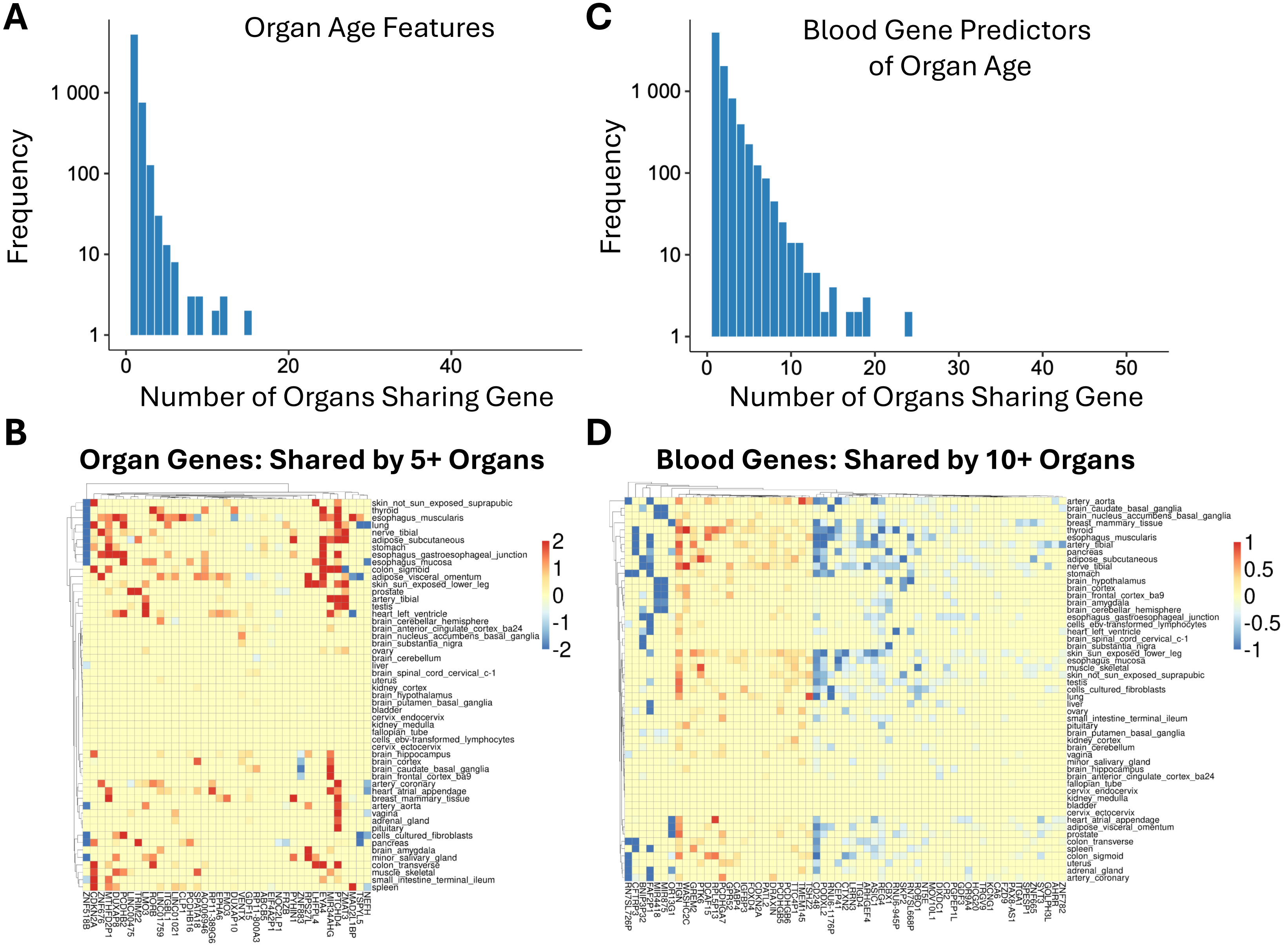

A
Organ Age Features
Frequency
1 000
100
10
1
0
20
40
Number of Organs Sharing Gene
C
Blood Gene Predictors of Organ Age
Frequency
1 000
100
10
1
0
10
20
30
40
50
Number of Organs Sharing Gene
B
Organ Genes: Shared by 5+ Organs
2
1
0
-1
-2
D
Blood Genes: Shared by 10+ Organs
1
0.5
0
-0.5
-1

**Supplementary Figure S2. Organ and blood genes in organ age models.** A) Organ genes shared in the organ age model constructed with genes correlated with age and predicative using blood. B) Coefficients of genes shared by five or more organ models. C) Blood genes shared in the blood-organ age models constructed from blood expression correlated with organ age. D) Coefficients of genes shared by ten or more blood-organ models.

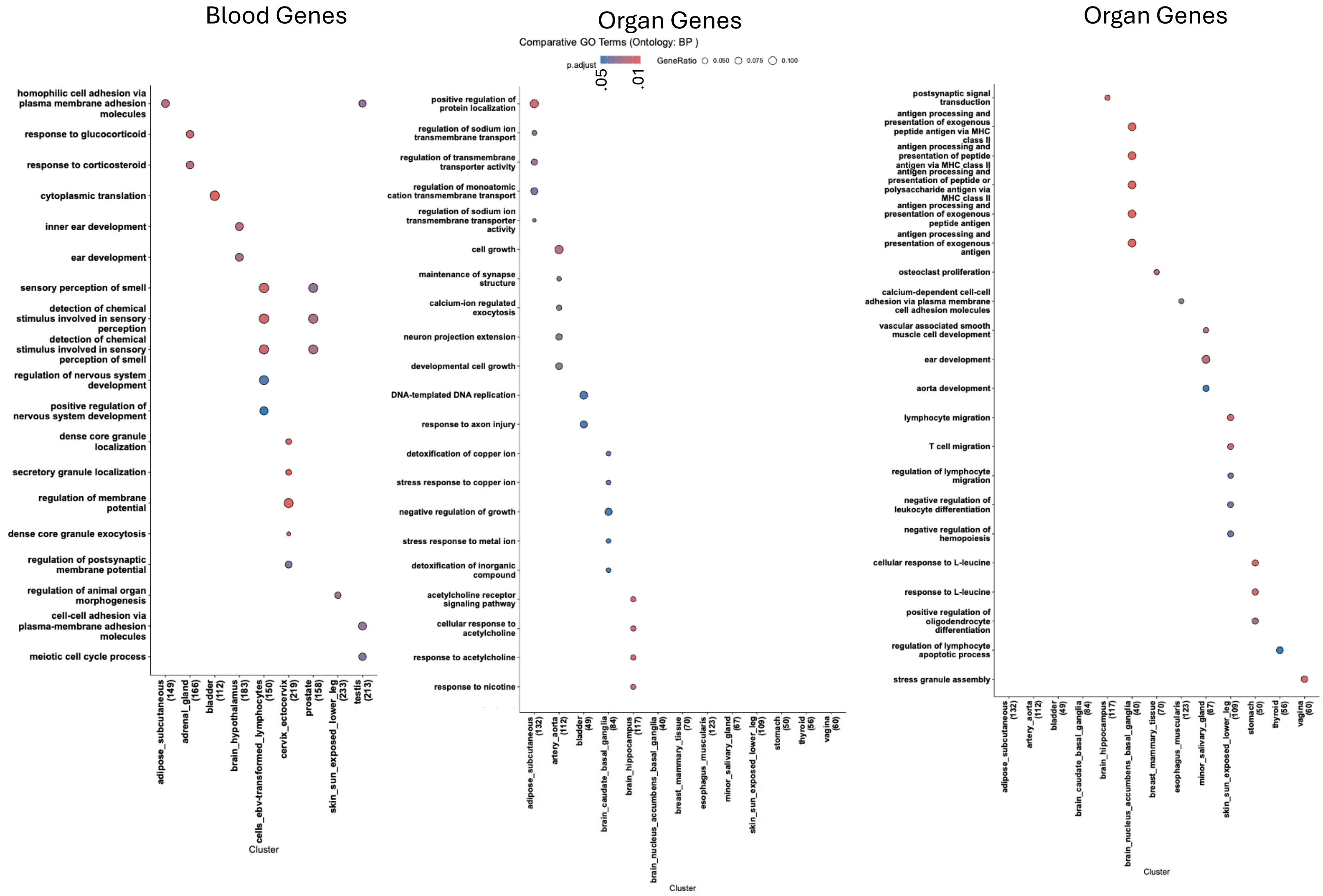


**Supplementary Figure S3. Biological process pathway enrichment of organ genes from organ age models and blood genes from blood-organ age models.** Enrichment of gene ontology biological process pathways were not shared between organs using organ genes. Fewer organs were enriched for pathways with blood genes.

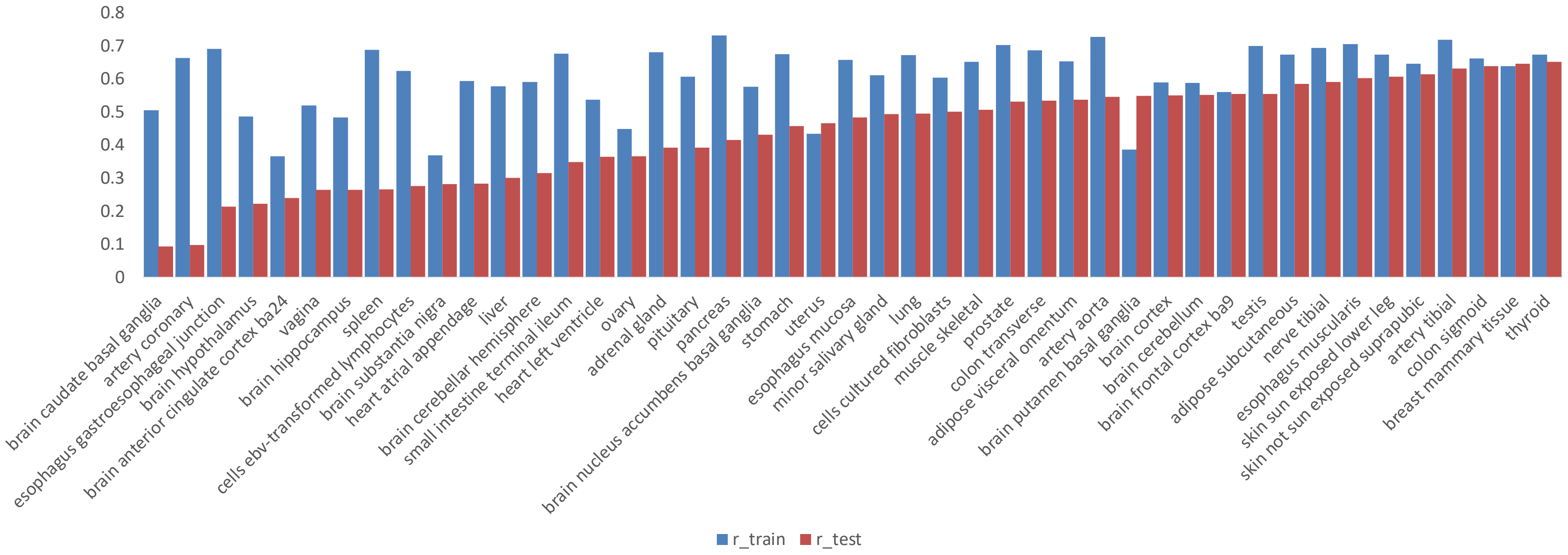


**Supplementary Figure S4. Pearson correlation of training and testing blood-organ age models.** Pearson correlation for each model was calculated in the training and testing set and sorted by the testing set. The testing set has lower correlation with age than the training set with most models.

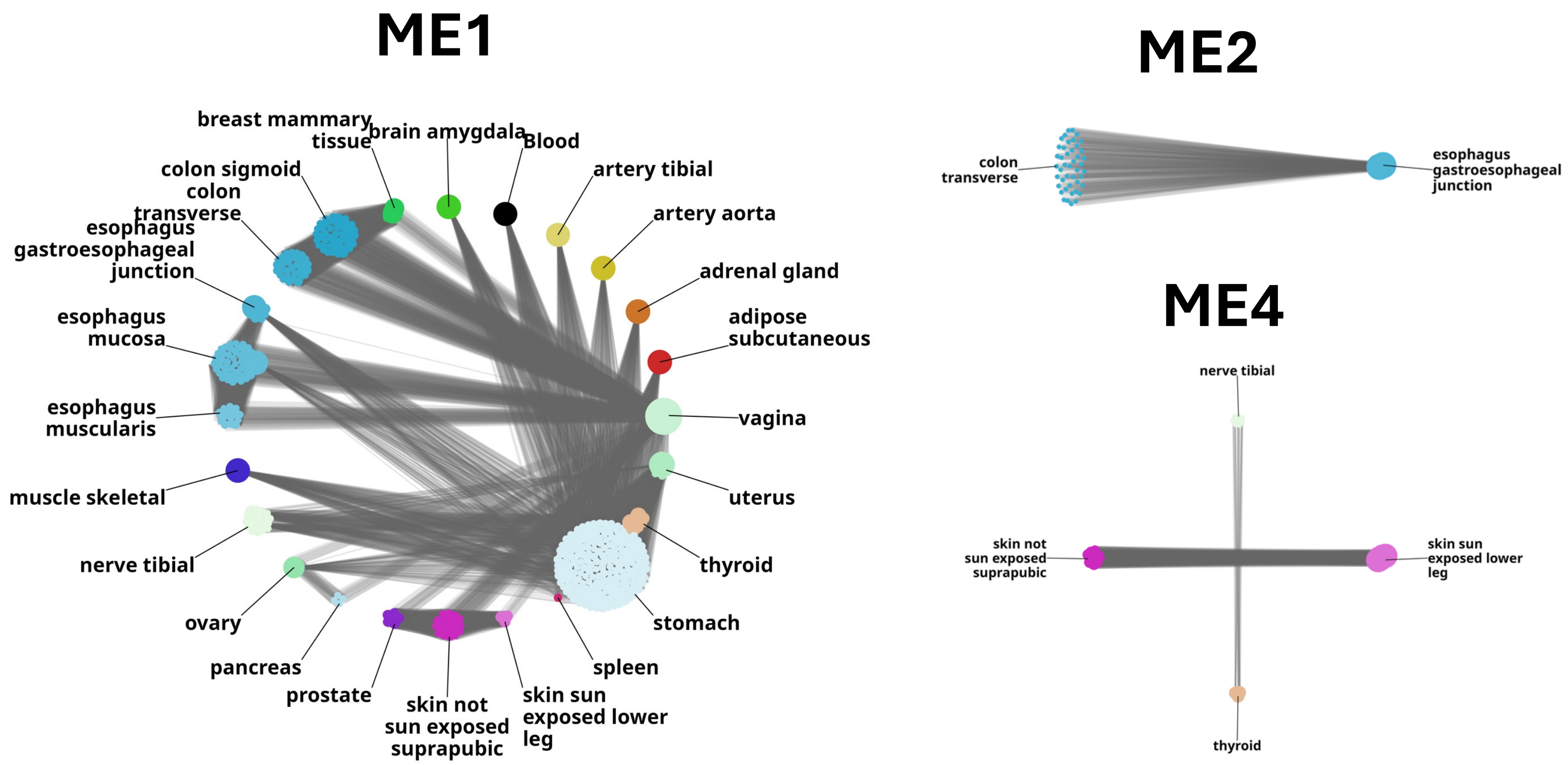


**Supplementary Figure S5. Whole genome multi-organ correlation analysis.** The most connected inter-organ edges are shown for each module significantly associated with age. Node sizes is calculated from the number of edges and edge width drawn based on the TOM matrix values. ME2 and ME4 were the only other two of 31 modules that contained more than one organ system.

### IMDB Mortality Predicitions from Image-Organ-Age

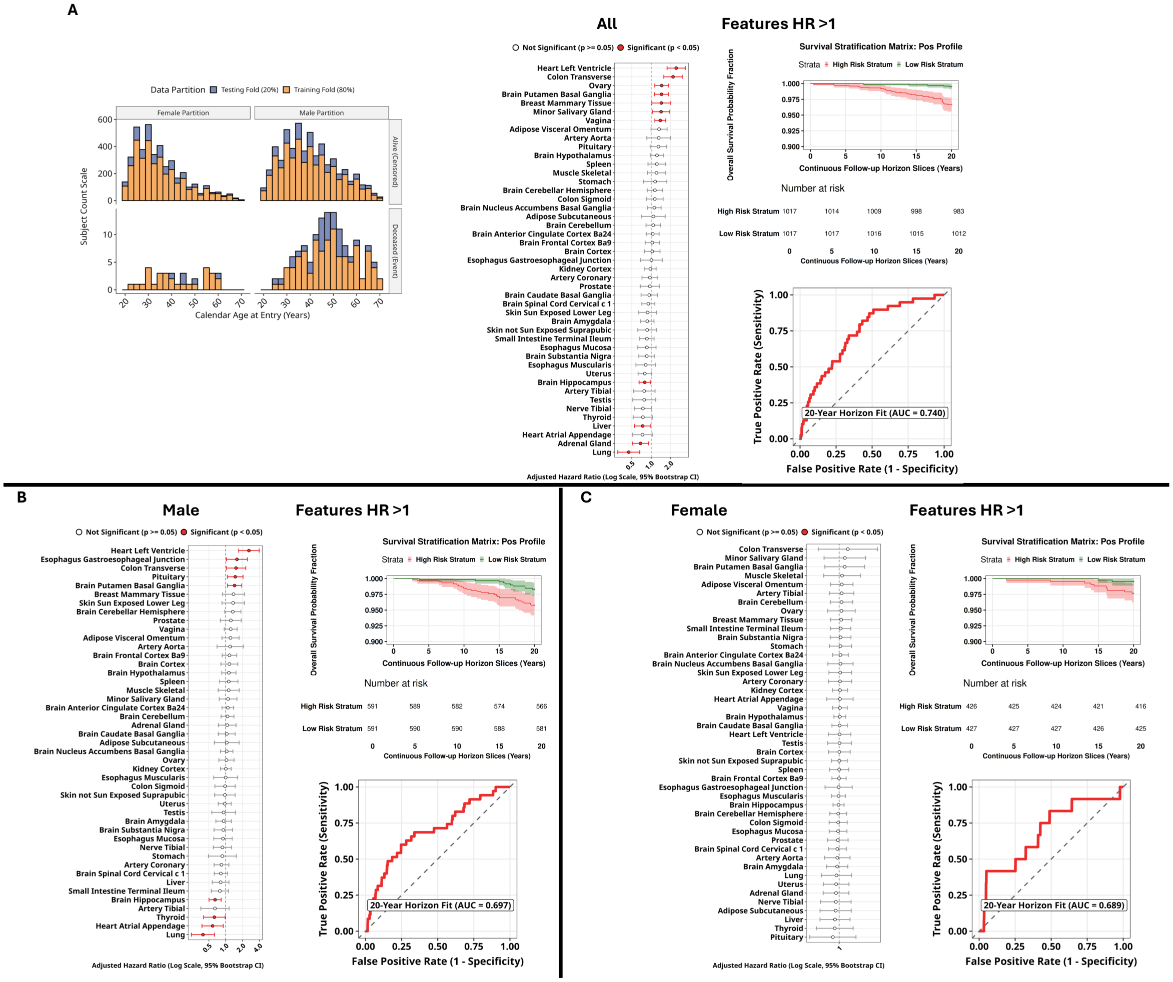


**Supplementary Figure S6. Facial-organ-age and mortality prediction in the IMDB-WIKI test set.** 1A) Groups were split into 80% training and 20% testing with models build on training data and survival plots and AUC calculated from test set. 1B) Multivariate cox analysis identified nominally significant increase and decreases in facial-organ-age scores associated with mortality. 1C) Survival plot splitting individuals by risk calculated as the median predicted value from features with HR >1 in multivariate cox partial hazard analysis. The ROC curve of the test set to correctly rank a mortality event has an AUC of 0.74. When adjusting for the confounding effects of each organ age, some organ age scores show protection as they increase. These nuanced features are more difficult to interpret and not included in the final model. 2) Samples were split into male only and used to define risk and map survival plots. 3) Samples were split into female only and used to define risk and map survival plots.

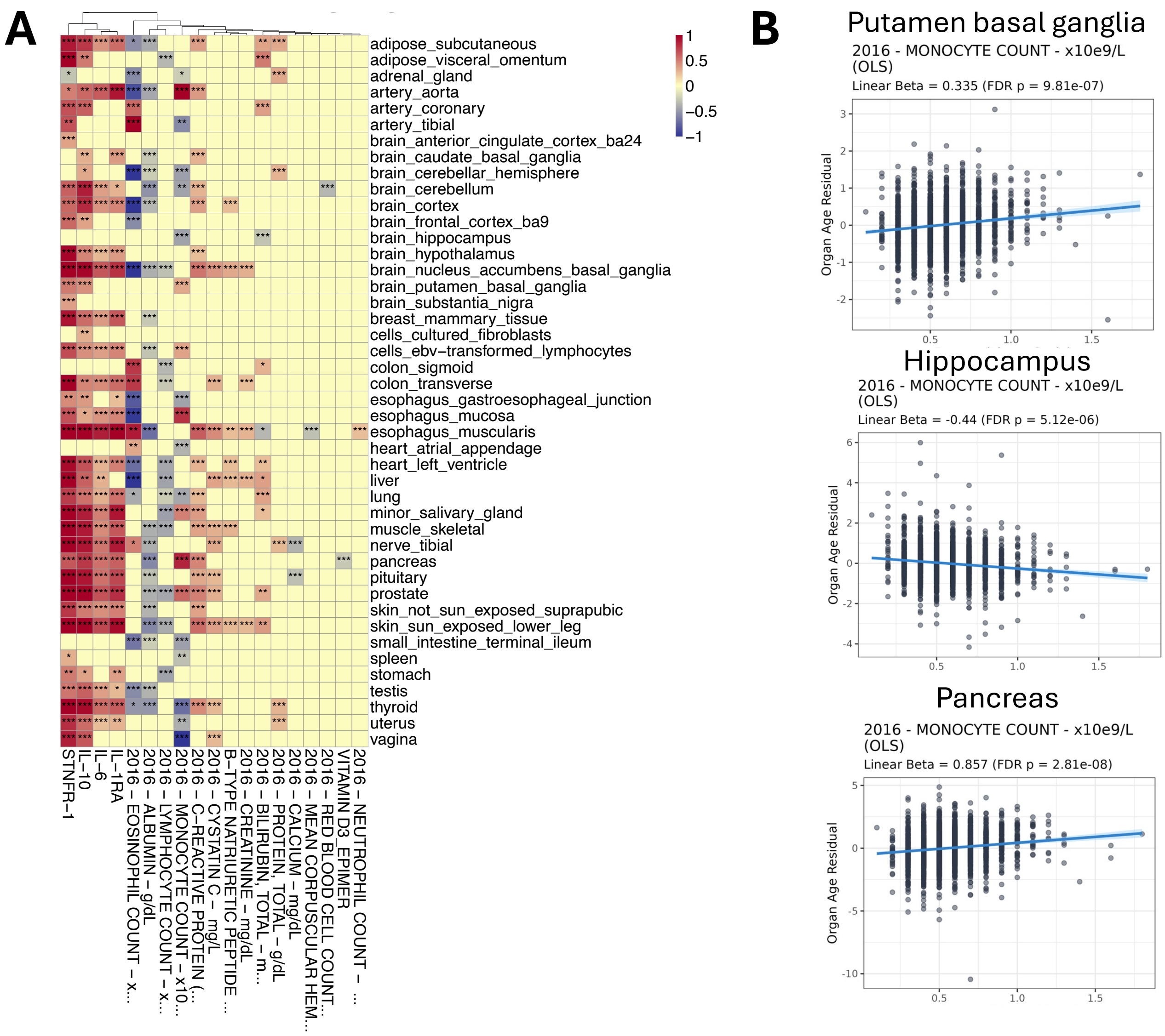


**Supplementary Figure S7. Blood-organ age associations with clinical and cognitive features in HRS.** A) The beta estimates between organ age residuals and clinical features where at least one feature had one organ with a significant correlation is shown. B) Monocyte counts for three of the organs age residuals.

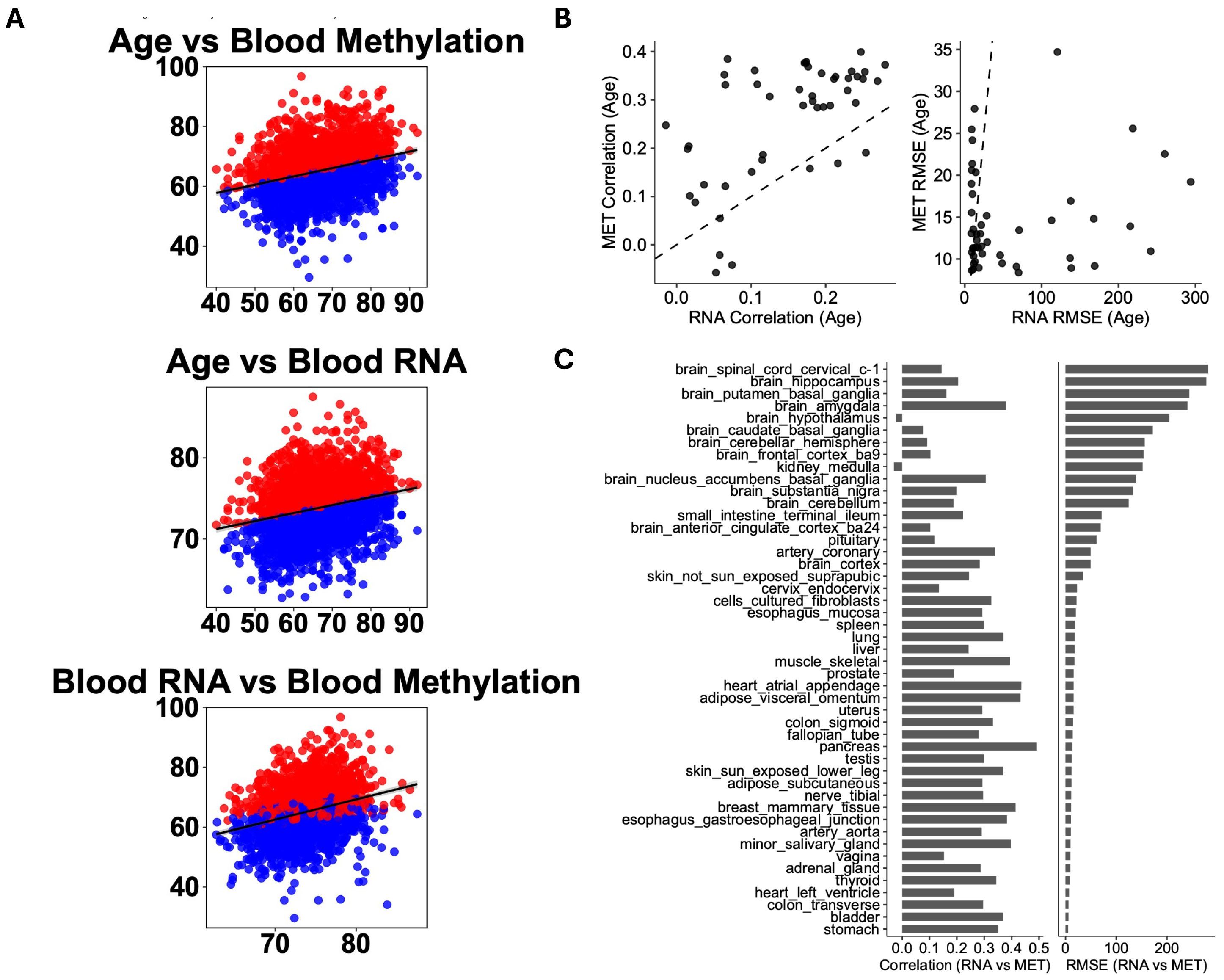


**Supplementary Figure S8. Blood-organ age and methylation-blood-organ age in FHS test dataset.** A) Scatter plots for adipose subcutaneous organ for methylation-blood-organ age versus chronological age, blood-organ age versus chronological age and blood-organ age versus methylation-blood-organ age. B) Comparison of Pearson correlation between blood-organ age and age (x-axis) and methylation-organ-age and age (y-axis) on the left and RMSE on the right. With FHS methylation-organ-age has higher correlation and lower RMSE with age than blood-organ age. This is likely due to the technical differences between assay types as FHS is microarray based. C) Change in correlation and RMSE between blood-organ age and methylation organ age for each organ.

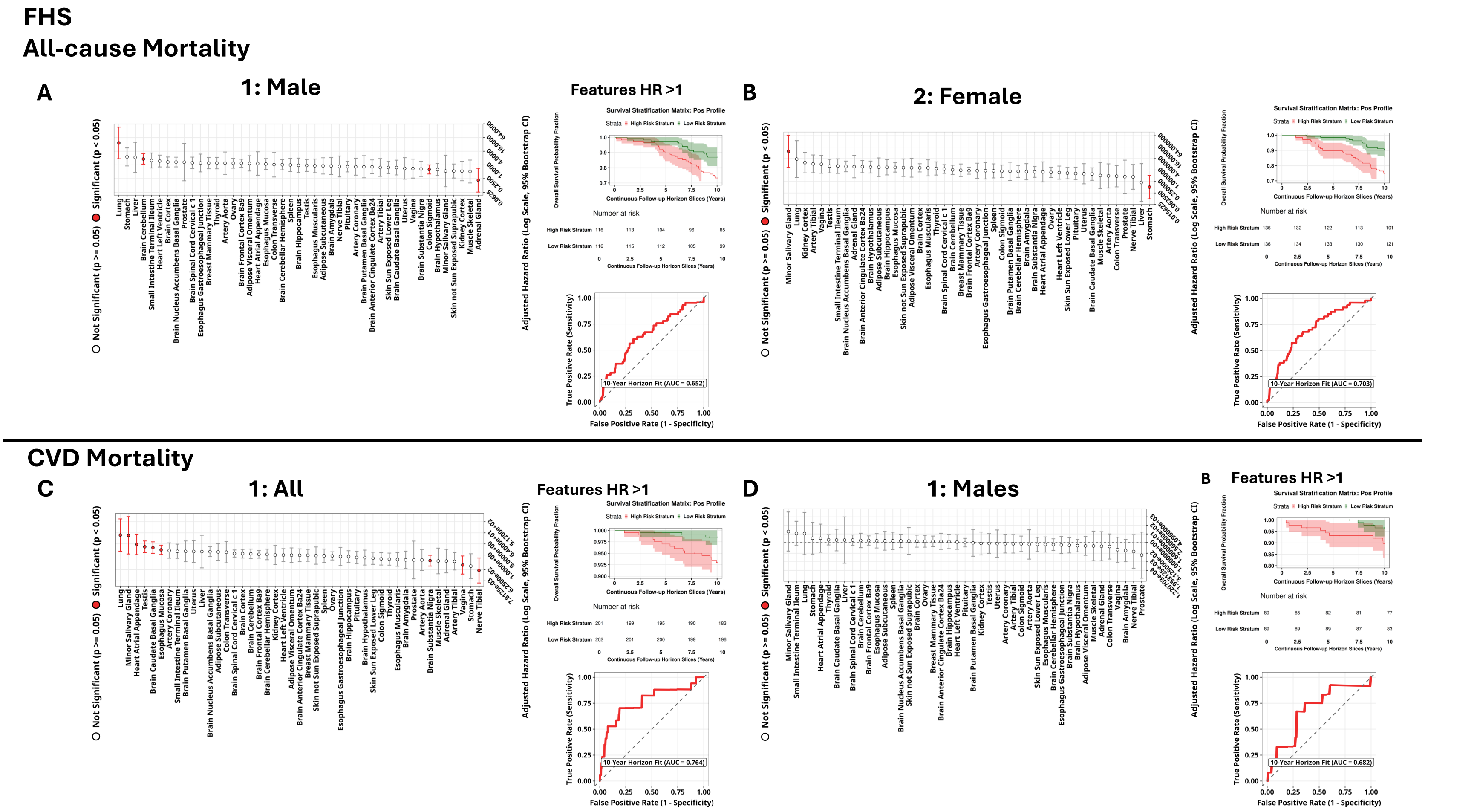


**Supplementary Figure S9. Methylation-blood-organ age predicts all-cause mortality in male and females and cardiovascular mortality in all and male individuals in FHS.** A) Males in FHS were used to create a male-specific all-cause mortality model. B) Females in FHS were used to create a female-specific all-cause mortality model. A) All individuals in FHS were used to create a cardiovascular mortality model. B) Males in FHS were used to create a male-specific cardiovascular mortality model.

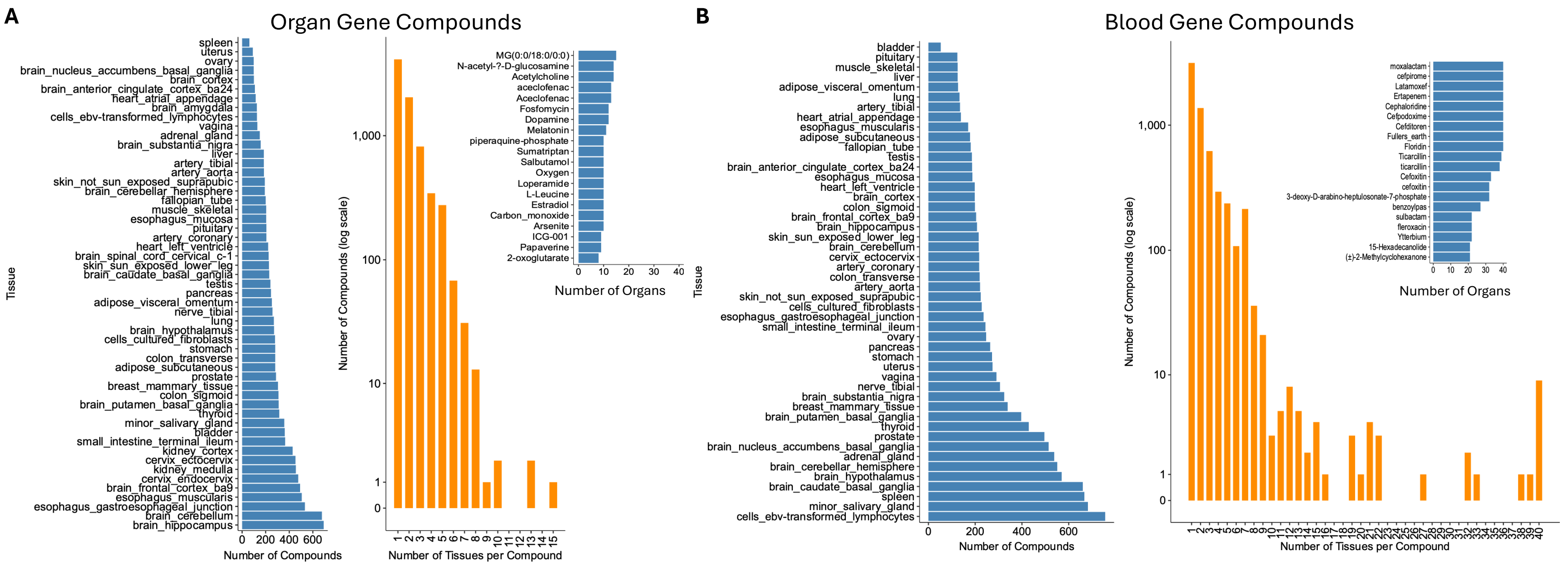


**Supplementary Figure S10. Compounds with translational potential to improve organ age and blood-organ age.** A) Drugs and food compounds from three databases were analyzed for enrichment with genes from the organ age models by gene compound enrichment analysis (GCEA). Some organs had up to 600 potential compounds that may ameliorate organ age. One monoacylglycerol compound targets genes from up to 15 different organs. B) Drugs and food compounds from three databases were analyzed for enrichment with genes from the blood-organ age models. As blood genes are more commonly shared between organs these compounds were more likely to associate with one than more organ.
